# Structural insights into the mechanism of the human SGLT2–MAP17 glucose transporter

**DOI:** 10.1101/2023.01.23.524988

**Authors:** Masahiro Hiraizumi, Tomoya Akashi, Kouta Murasaki, Hiroyuki Kishida, Taichi Kumanomidou, Nao Torimoto, Osamu Nureki, Ikuko Miyaguchi

## Abstract

Selective sodium-glucose cotransporter 2 (SGLT2) plays an important role in glucose reabsorption. SGLT2 inhibitors suppress glucose reabsorption from the kidneys, thus reducing blood glucose levels in type 2 diabetes patients. We and other groups have developed several SGLT2 inhibitors starting from a natural product, phlorizin, but their action mechanisms remain unknown. Here, we elucidated the physiological hSGLT2–MAP17 complex structures bound to five SGLT2 inhibitors using single-particle cryo-electron microscopy. Canagliflozin, dapagliflozin, TA-1887, and sotagliflozin were bound in the outward-facing structure, whereas phlorizin was bound in the inward-open structure. The phlorizin–hSGLT2 interaction biochemically exhibited biphasic binding. Phlorizin weakly binds, via the phloretin motif, from its intracellular side near the Na^+^-binding site, while strongly interacts from its extracellular side. Unexpectedly, bound Na^+^ stabilizes the outward-open conformation, while its release allows the transporter to adopt inward-open state. Our results first visualized the Na^+^-binding and inward-open conformation of hSGLT2–MAP17, clarifying the unprecedented Na^+^-dependent sugar transport mechanism with MAP17 acting as a scaffold, and may pave the way for development of next-generation SGLT inhibitors.

## Introduction

Type 2 diabetes mellitus is characterized by persistent hyperglycemia caused by inadequate insulin action. Chronic high blood sugar level damages blood vessels, causing serious health problems, such as nephropathy and cardiovascular disease. The primary treatment for diabetes is blood glucose control; drugs such as selective sodium-glucose cotransporter (SGLT1 and 2; also known as SLC5A1 and 2) inhibitors^1^ hold great promise for reducing blood glucose levels. Human SGLT1 and SGLT2 are responsible for the reabsorption of plasma glucose in the proximal tubules after filtration through the renal capillaries^2^. SGLT2 is located in the S1 and S2 segments of the proximal tubule and absorbs 90% of plasma glucose; SGLT1, which has higher glucose affinity, is located in the S3 segment of the proximal tubule and absorbs the remaining 10%^3^.

SGLT2, a transmembrane protein with 14 helices (Fig. 1a) and expressed specifically in the kidneys, has 60% sequence homology to SGLT1, which is expressed in the small intestine as well. SGLT2 inhibitors that suppress glucose reabsorption and promote urinary excretion are considered a promising therapeutic tool to manage blood glucose levels in type 2 diabetes patients. SGLT2 inhibitor development initially focused on the natural product phlorizin (Figs 3b)^4^, an O-glucoside that is hydrolyzed by β-glucosidase in the intestine, making oral administration difficult owing to its poor metabolic stability. This was replaced by N- or C-glucosides, with better metabolic stability. C-glucosides, including canagliflozin, dapagliflozin, and empagliflozin, are marketed as approved drugs in USA, Japan and many countries (Figs 1b, 2g, h)^5^. SGLT2 inhibitors have ushered in a new phase of diabetes treatment, providing many benefits including, reduced risk of heart failure and kidney protection^6–9^. It was initially thought that rare mutations in the Na^+^–glucose cotransporter gene *SLC5A1* can cause lethal glucose–galactose malabsorption and that inhibitors with high specificity for SGLT2 over SGLT1 were necessary to treat diabetes. However, SGLT1 inhibition in the gastrointestinal tract leads to postprandial glucose excursion control and gastrointestinal hormone secretion^10^. Therefore, SGLT1 and SGLT2 dual inhibitors (e.g., sotagliflozin and LX-2761; Fig. 2e) are currently being developed for the treatment of diabetes^9,11^.

**Figure 1.**
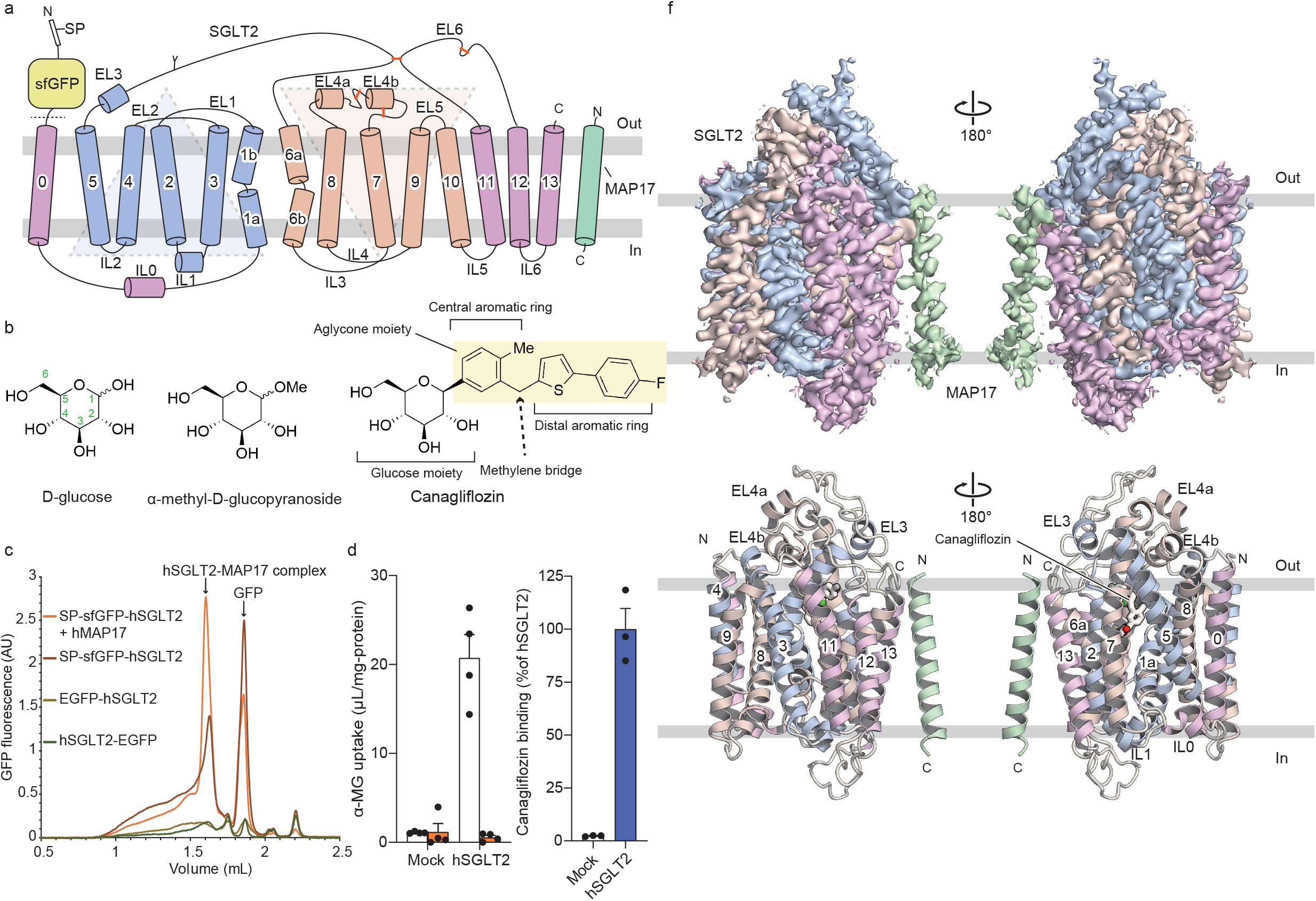
Biochemical and cryo-EM studies of the human SGLT2–MAP17 heterodimer. **a** Topological diagram of the hSGLT2–MAP17 heterodimer. Inverted repeats (IRs) are represented as light blue and light brown. The Y-shape indicates an N-glycosylation site. Disulfide bonds are shown as orange sticks. A signal peptide (SP) and super-folder (sf) GFP were fused to the N-terminus of hSGLT2. The dotted line indicates the site of protease activity. **b** Chemical structures of the substrates and a representative gliflozin. **c**, FSEC profiles of various types of GFP-tagged hSGLT2 without MAP17. Arrows indicate elution positions of the hSGLT2–MAP17 heterodimer and free GFP (GFP). **d** α-MG uptake in hSGLT2 and MAP17 expressing cells in the absence (white) or presence (orange) of 500 nM canagliflozin. Each column represents mean ± SEM (n = 4, biological replicates). **e** Canagliflozin (30 nM) binding assay for the hSGLT2- and MAP17-expressing cell membrane fraction. Each column represents mean ± SEM (n = 3, technical replicates). **f**, Overall structure of the human SGLT2-MAP17 complex. Cryo-EM maps (top) and ribbon models (bottom). The same color scheme was used throughout the manuscript, except for Fig. 4.

**Figure 2.**
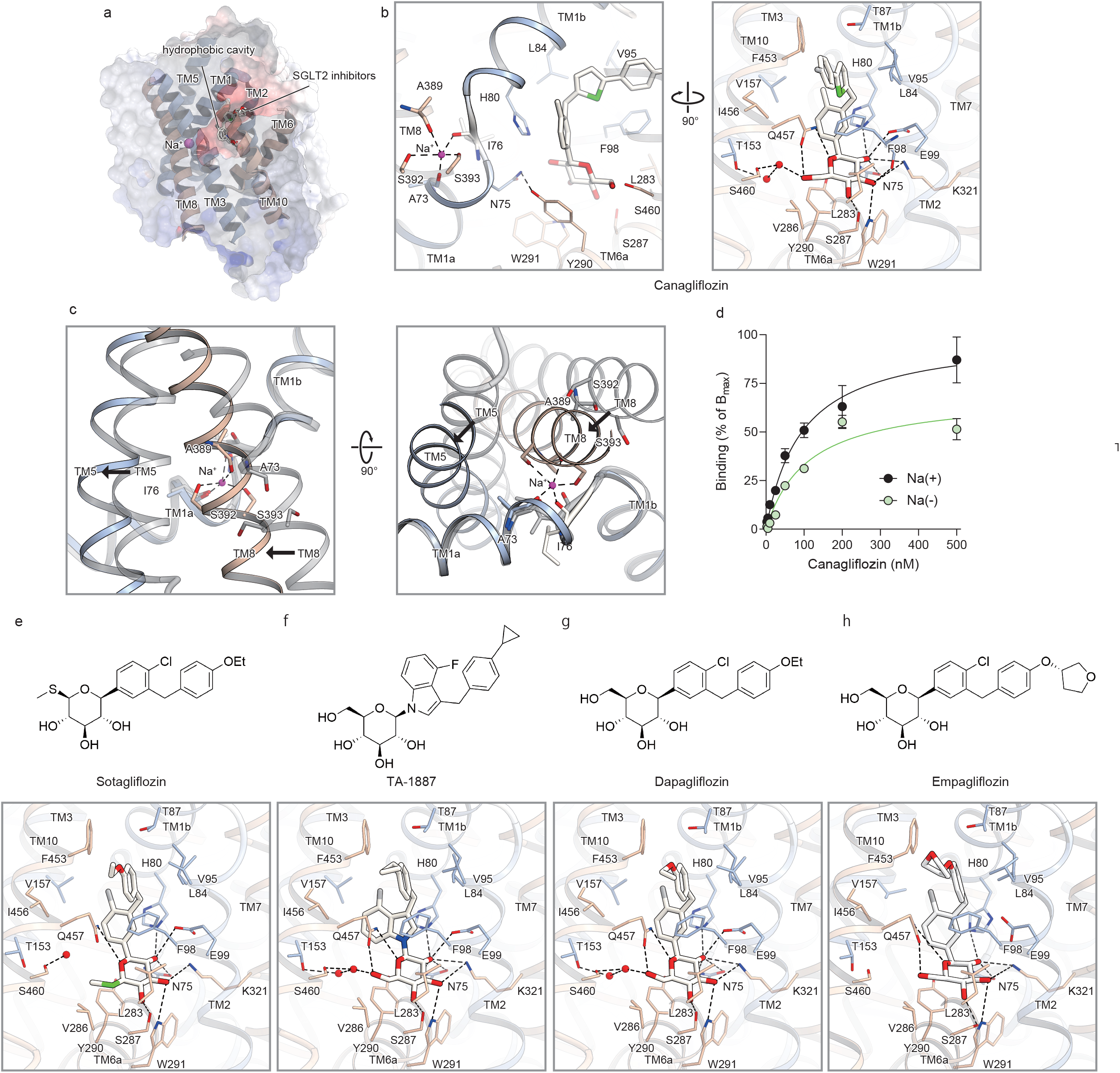
SGLT2-inhibitor-binding site of the outward-facing conformation. **a,** Cross-sections of the electrostatic surface potentials at the SGLT2 inhibitor-binding site. The potentials were displayed as a color gradient from red (negative) to blue (positive). **b**, Interactions between canagliflozin and hSGLT2. Canagliflozin and its interacting residues are shown as sticks. The hydrogen bonds are indicated by the black dashed lines. **c**, Conformation change from outward-opening (color) to inward-opening (grey) around the Na2 binding site. View from the lateral side of the plasma membrane (left) and from the cytoplasmic side (right). **d**, Concentration-dependent binding evaluation of canagliflozin in the presence and absence of Na^+^. Each point represents the mean ± SEM (n = 3, technical replicates). **e–h**, Chemical structures of inhibitors and their interactions with hSGLT2: sotagliflozin (**e**), TA-1887 (**f**), and dapagliflozin (**g**) in this study, and of empagliflozin (**h**) (previously reported; ^19^).

**Figure 3.**
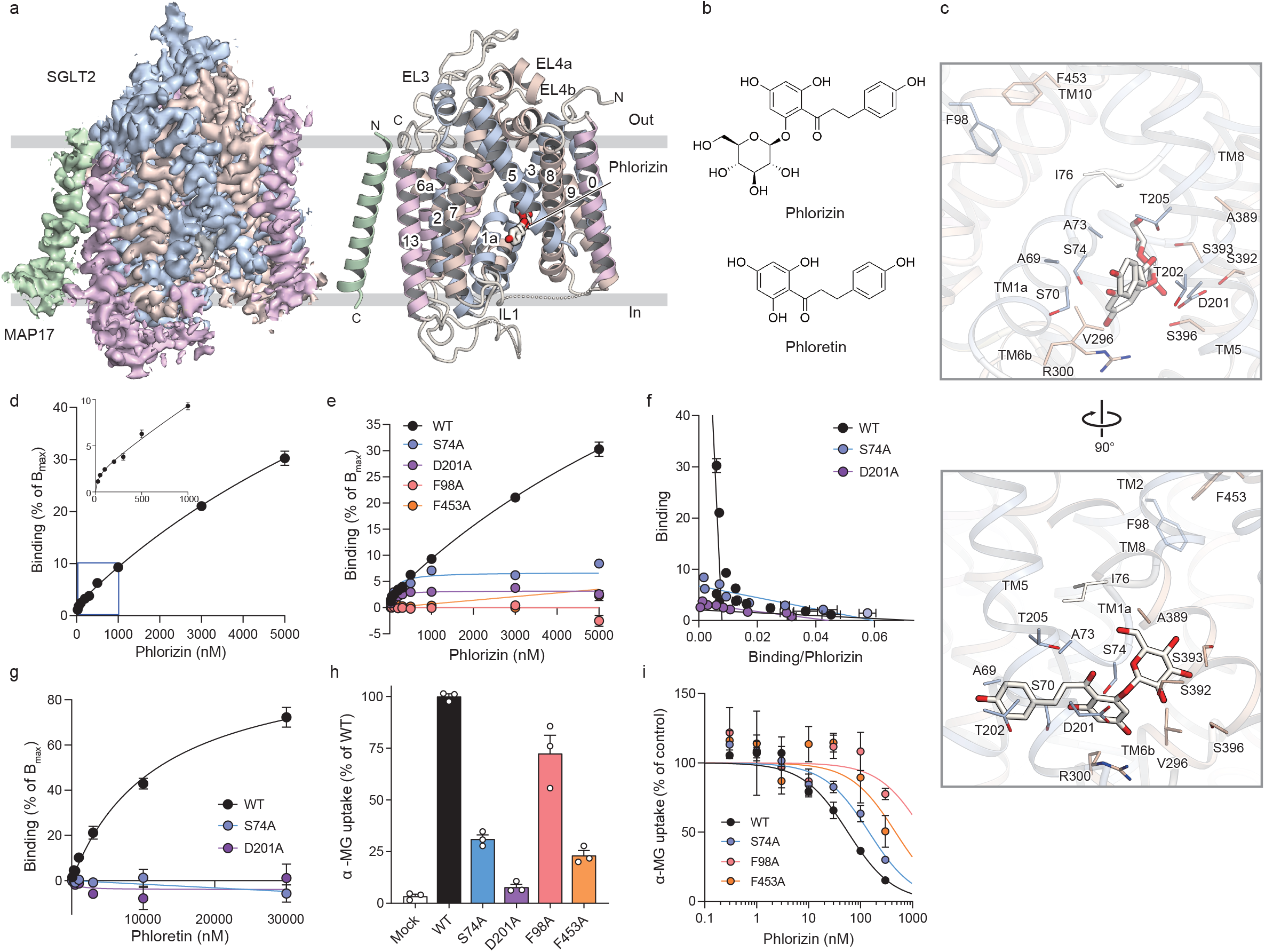
Phlorizin binding site of the inward-open conformation. **a**, The inward-open conformation of the hSGLT2–MAP17 complex. Cryo-EM maps (left) and ribbon models (right). **b,** Chemical structures of phlorizin and phloretin. **c**, Interactions between phlorizin and hSGLT2. **d**, Concentration-dependent binding of phlorizin to the wild-type hSGLT2. Each point represents the mean ± SEM (n = 3, technical replicates). **e**, Concentration-dependent binding of phlorizin in wild-type and mutant hSGLT2. Each point represents the mean ± SEM (n = 3, technical replicates). **f,** Eadie-Hofstee plot analysis of phlorizin binding in wild-type and mutant hSGLT2. Each point represents the mean ± SEM (n = 3, technical replicates). **g**, Concentration-dependent binding of phloretin in wild-type and mutant hSGLT2. Each point represents the mean ± SEM (n = 3, technical replicates). **h**, Uptake assay of α-MG in wild-type and mutant hSGLT2. Each column represents mean ± SEM (n = 4, biological replicates). **i**, Inhibitory effect of phlorizin on α-MG uptake by wild-type and mutant hSGLT2. Each point represents the mean ± SEM (n = 4, biological replicates).

**Figure 4.**
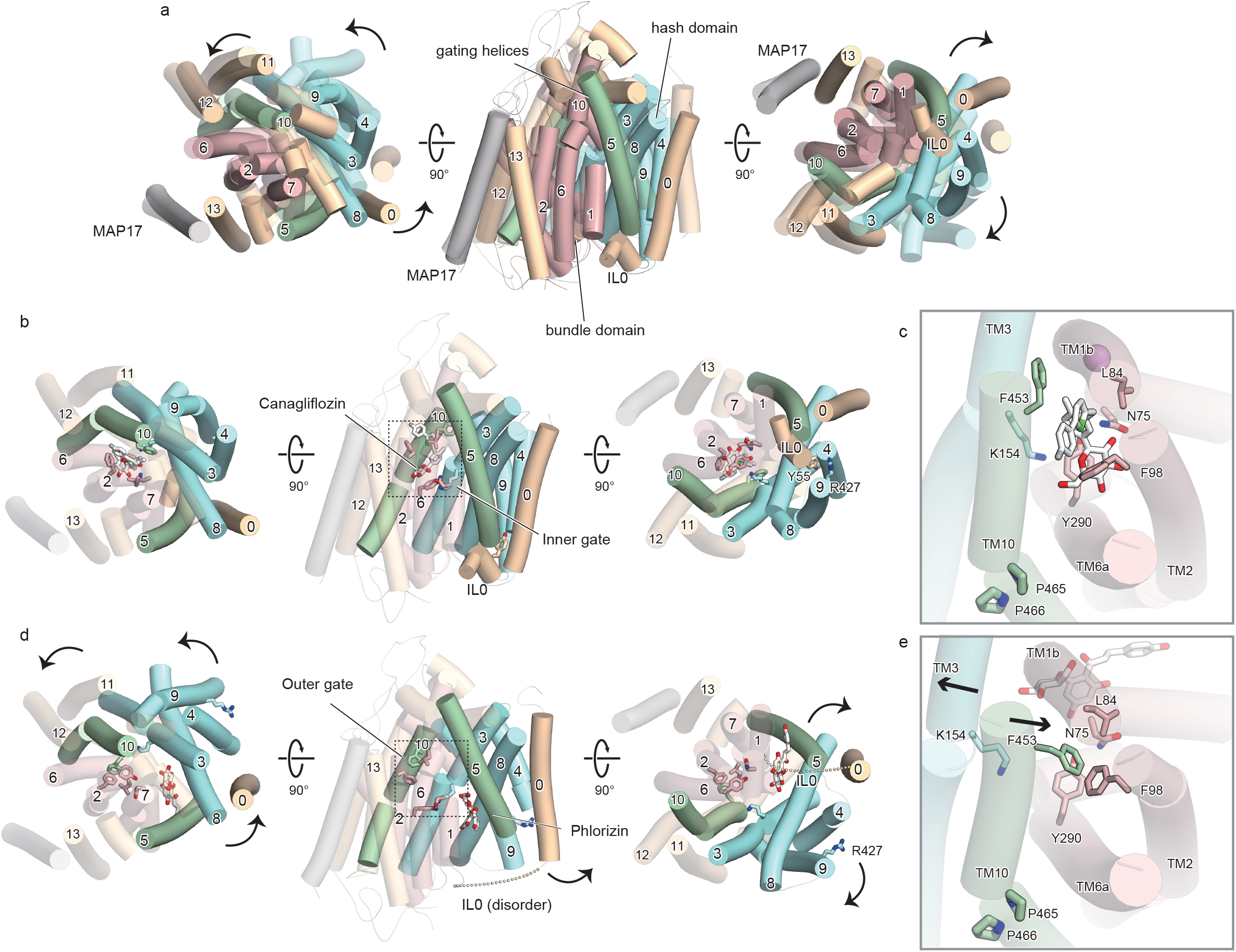
Comparison of the outward-facing and inward-open conformations of hSGLT2. **a**, The outward-facing (colored) and inward-open (transparent) structures when their bundle domains (TM1, −2, −6, and −7; red) are superimposed. MAP17 (grey) and TM13 (light orange) overlap well between the conformations, but the hash domain (TM3, −4, −8, −9; blue), gate helices (TM5, −10; green), TM0, −11 and −12 (light orange) are in the inward-opening conformation. **b**, The outward-facing conformation of the hSGLT2–MAP17 complex with canagliflozin viewed from the exoplasm (left), side (center), and cytoplasm (right). **c**, Substrate sugar-binding site and external vestibule of the outward-facing conformation from the exoplasm. **d**, The inward-open conformation of the hSGLT2–MAP17 complex with phlorizin viewed from the exoplasm (left), side (center), and cytoplasm (right). **e**, Substrate sugar-binding site and external vestibule of the inward-open conformation from the exoplasm.

SGLT1 and SGLT2 belong to the LeuT transporter family and are conserved in all bacterial and animal taxa, with six isoforms in humans^12,13^. Structural homology modeling studies have been conducted using SGLT from *Vibrio parahaemolyticus* (vSGLT) and similar protein structures, such as SiaT^14,15^, because of the difficulties in protein preparation and structural analysis. A functional model of SGLT2 glucose uptake has been proposed: SGLT2 undergoes a conformational change to an outward-facing conformation by binding to a single Na^+^ ion (at Na2 site^16^) before binding to the substrate, depending on Na^+^ concentration gradient across the plasma membrane, which then promotes glucose binding. Subsequently, Na^+^ and sugars are incorporated into the cell in an inward-open conformation^3,12,13^. In contrast, SGLT1 requires two Na^+^ ions (at Na2 and Na3 sites) for glucose transport, which probably affects its glucose affinity^16^. The structural understanding of the human SGLT family was advanced by the cryo-electron microscopy (cryo-EM) structures of SGLT1 and SGLT2. The structure of the hSGLT1 apo-form with consensus stabilizing mutations and molecular dynamics calculations, revealed the mechanism of glucose binding and selectivity and water permeability^17^. The activity of hSGLT2, whose gene has long been difficult to clone, is greatly enhanced by the co-expression of MAP17 (PDZK1P1), an essential auxiliary subunit of hSGLT2^18^. Determination of the hSGLT2 structure via MAP17 tethering and the introduction of several mutations revealed that it is bound by empagliflozin in an outward-facing conformation^19^. Furthermore, introduction of MAP17 tethering and several mutations in hSGLT1 revealed that the dual inhibitor LX2761 was bound in the outward conformation of hSGLT1^20^. The role of amino acid residues in the binding of SGLT inhibitors in these two outward conformations was discussed. However, the function of Na^+^ remains unknown, because these previous studies did not detect Na^+^, which should bind to hSGLT before binding to the substrate. Regarding the binding mode of inhibitors, only C-glucoside inhibitors have been shown to bind in the outward conformation, while the binding mode of O- and N-glucoside type inhibitors, including phlorizin, still needs to be elucidated. Moreover, the conformational changes of SGLT2 between the outward and inward conformations as well as the role of MAP17 as a scaffold have yet been unclarified.

Here, we performed cryo-EM single-particle analyses to determine the structure of the genuine hSGLT2–MAP17 complexes with five inhibitors (canagliflozin, dapagliflozin, TA-1887, sotagliflozin, and phlorizin). This characterization of Na^+^-binding outward-facing structures and inward-open structures with MAP17 as a scaffold, together with transport and binding assays, significantly clarifies the molecular features of hSGLT2–MAP17, SGLT2 inhibition, and sugar transport, which helps achievement of glycemic control in patients.

## Results and discussion

### Structural determination of the hSGLT2–MAP17 heterodimer

We performed cryo-EM analysis of hSGLT2 to elucidate its inhibitory mechanism. To obtain a stable and homogeneous sample, we first examined, by fluorescence-detection size-exclusion chromatography (FSEC)^21^, the expression of hSGLT2 whose N or C terminus was fused with enhanced green fluorescent protein (EGFP). However, we were unable to detect hSGLT2 expression (Fig. 1c); given that its N-terminus is exposed to the extracellular side, we hypothesized that N-terminal fusion of a signal sequence and super folder GFP (sfGFP) would improve the protein expression^22^. Fusing the human trypsinogen 1-derived signal peptide and sfGFP to the hSGLT2 N-terminus revealed a peak, indicating that hSGLT2 can be solubilized by detergents (Fig. 1c). Co-expression of MAP17 and hSGLT2 caused a high molecular-weight shift in FSEC, suggesting heterodimer-complex formation. LC-MS/MS confirmed that signal peptides and sfGFP fused with hSGLT2–MAP17-expressing cells could take up α-methyl-D-glucopyranoside (α-MG; Fig. 1b); this uptake was sensitive to canagliflozin (Fig. 1d). We used LC-MS/MS to determine whether the prepared membrane fractions maintained their inhibitor-binding activity, verifying the binding of multiple inhibitors to hSGLT2 (Fig. 1e; Supplementary Fig. 1c). Membrane fraction of hSGLT2–MAP17-expressing cells was solubilized with N-dodecyl β-d-maltoside (DDM) micelles in the presence of SGLT2 inhibitors, purified using GFP nanobody-affinity chromatography, followed by cleavage of sfGFP using protease, and subjected to gel-filtration column chromatography (Supplementary Fig. 1a, b).

Purified hSGLT2–MAP17 complexes were subjected to cryo-EM single-particle analyses with five inhibitors (canagliflozin, dapagliflozin, TA-1887, sotagliflozin, and phlorizin; Supplementary Figs 3–6, 8). The acquired movies were motion-corrected and processed in RELION^23,24^ providing cryo-EM maps at overall resolutions of 2.6–3.3 Å, according to the gold-standard Fourier shell correlation (FSC) = 0.143 criterion (Supplementary Data Table 1). All of the potential maps contained disordered regions but were sufficient for building structural models of proteins and inhibitors (Supplementary Figs 7, 9). The overall structure exhibits a LeuT fold comprising 14 membrane-spanning helices (TM0–TM13). Helices TM1–5 and TM6–10 formed an inverted repeating structure (Fig. 1a). N-glycan, attached to N250 of hSGLT2, was identified (Supplementary Figs 7, 9). Canagliflozin, dapagliflozin, sotagliflozin, and TA-1887 were found to bind to the outward-facing conformation, whereas phlorizin was found to bind to the inward-open conformation. Because of their high flexibility, the N-terminal loop of hSGLT2 (15–20 amino acids), the IL6 loop between TM12 and TM13, and the extracellular or intracellular region of MAP17, were not visible in either structure. The inward-open structure does not exhibit density for IL0 between TM0 and TM1. In MAP17, a single transmembrane helix interacts with TM13 of hSGLT2, consistent with the published interaction between MAP17 and hSGLT2^19^. The extracellular half interacts closely with hydrophobic residues and lipids, while the intracellular half does not interact with hSGLT2 (Fig. 1f; Supplementary Fig. 10).

### The inhibitor-bound hSGLT2 structure and the roles of the Na2 site

The C-glucoside inhibitors (canagliflozin, dapagliflozin, and sotagliflozin), and the N-glucoside TA-1887, are bound to the central hydrophobic cavity of the topologically inverted repeats (IRs) (TM1, TM2, TM3, TM6, and TM10; Figs 1a, 2a) of hSGLT2 in outward-facing conformation. This cavity was negatively charged, favoring the binding of positively charged ions such as Na^+^. The binding mode was the same as that in the recently reported SGLT2-empagliflozin structure^19^; for the main chain, with root-mean-square deviation (RMSD) ranging from 0.66 to 0.85 Å.

SGLT2 couples the transport of one Na^+^ ion and one glucose molecule. The Na^+^-binding Na2 site, which is conserved in many LeuT-fold transporters, was located near the middle bend of TM1 (Fig. 2b). In SGLT2, the Na2 site is thought to be formed by the backbone carbonyls of A73, I76, and A389 and the side chain oxygens of S392 and S393 (Fig. 2b), based on the vSGLT^15^ and SiaT^25^ alignment (Supplementary Fig. 2). The density corresponding to Na^+^ at the Na2 site was confirmed in all four outward-facing structures studied here (Supplementary Fig. 11). As predicted from the alignment, Na^+^ interacts with A73 and I76 of TM1 and with A389, S392, and S393 of TM8, resulting in a trigonal bipyramidal form (Fig. 2b). Thus, TM1 and TM8 are connected by Na^+^ and form part of the substrate-binding site. In the outward-facing structure, this connection not only brought TM1 and TM8 closer together but also allowed the entire outer region of the glucose-binding site, such as TM5, to move outward, causing the inward-open to outward-facing conformational change (Fig. 2c). The binding affinity of canagliflozin to hSGLT2 was lower in the Na^+^ absent conditions (Fig. 2d, Supplementary Data Table 2). Therefore, Na^+^ binds to hSGLT2 before the substrate or inhibitors binding, allosterically stabilizing the outward-facing structure. Our current findings first clarified the role of sodium ions in the transport state exchange. This is consistent with the fact that, in SGLT1, substitutions of S392 and S393 at the Na2 site significantly reduced glucose uptake^26^.

### Implication of the role of Na3 site of SGLT1

SGLT1 requires two Na^+^ ions for sugar transport and has an Na3 site in addition to the Na2 site^3^. In SGLT2, the region corresponding to SGLT1 Na3 site, located on the cytoplasmic side and away from the glucose-binding pocket, is occluded by the side chain carboxyl group of D201 in TM5 and the backbone carbonyls of S392, S393, A395, and S396 in TM8 (Fig. 3c and Sup. Fig. 2 and 11b). In SGLT1, the residue corresponding to A395 in hSGLT2 is replaced by Thr, which probably contributes to Na^+^ binding^17^. This is consistent with the fact that no density corresponding to Na^+^ was observed at the Na3 site in the present hSGLT2 structures (Supplementary Fig. 11b).

It has been proposed that, as at the Na2 site, Na^+^ at the Na3 site binds before substrate binding and is taken up by the cell before the substrate^14^. In the present outward-facing structure of hSGLT2, the region corresponding to SGLT1 Na3 site is located very close to Na2 site so as to partially share the same amino acid residues, suggesting that the Na3 site, in addition to Na2 site, of SGLT1 also stabilizes the outward-open, substrate-binding structure by connecting TM1 to TM5, as Na2 connects TM1 and TM8 in SGLT2. Taken together, this may contribute to the higher affinity of SGLT1 for its substrate^16^ but its slower turnover relative to SGLT2^27^.

### The hSGLT2 outward-facing conformation in complex with inhibitors

SGLT2 inhibitors were composed of glucose and aglycone moieties (Fig. 1b). The aglycone moiety comprises two aromatic rings that bend at the methylene bridge and extend toward the extracellular space (Fig. 2a, b). The glucose moiety of all gliflozin inhibitors stacks with the aromatic side chain of the inner gate, Y290. The hydroxyl groups of the glucose moiety form hydrogen bonds in the side chains of N75, H80, E99, S287, W291, K321, and Q457, and in the main chain carbonyl group of F98 (Fig. 2b); these residues were conserved in SGLT1 (Supplementary Fig. 2). A water molecule-like density was observed near S460 in the structures of canagliflozin, dapagliflozin, and TA-1887 (Fig. 2b and Supplementary Fig. 7). It has been reported that the corresponding T460 mutant of SGLT1 has reduced sugar transport activity^28^. Therefore, these S460-bound water molecules may also participate in D-glucose transport in hSGLT2.

The IC50 values of the five compounds for hSGLT2 were 1–6 nM, with little difference, while sotagliflozin, an SGLT1/2 dual inhibitor, had >10-fold stronger activity toward SGLT1 (IC50 of 40 nM) than the other inhibitors^6^. Sotagliflozin has a methylsulfanyl group at C5 of glucose, whereas dapagliflozin has a hydroxymethyl group which has a hydrogen bonding interaction with Gln457 and a water molecule-mediated interaction with S460 (Fig. 2e, g, Supplementary Fig. 7). In SGLT1, S460 of hSGLT2 near C5 was replaced by Thr, V286 by Leu, and L283 by Met, causing the inhibitor-subtype selectivity^19^.

In all of the gliflozin inhibitors, the central aromatic ring is exposed to the hydrophobic cavity formed by TM1, TM3, and TM10 (Figs. 2e-h): in TA-1887, this is a benzylindole ring, which extends into the hydrophobic cavity (Fig. 2f). The hydrophobic substituents at the para position affect hSGLT2 inhibitory activity^29,30^, and are all located at approximately the same position (Fig. 2a, b, and e–h). V157, located in TM3 in hSGLT2, is replaced by A160, smaller than Val, in hSGLT1. This site is potentially selective for hSGLT2, because the hydrophobic pocket sizes should be different. Molecular dynamics simulation frames in the hSGLT1 with mizagliflozin showed that A160 is important for the selectivity, which is consistent with the present results^31^.

Distal aromatic rings form a long hydrophobic aglycon tail that extends into the extracellular vestibule. In all gliflozins, each tail formed a T-shaped π-π stacking with F98 of TM2 and was surrounded by hydrophobic amino acids such as Leu84 of TM1, V95 of TM2, and F453 of TM10 (Fig. 2b, e–h). Canagliflozin extends with fluorophenyl via a thiophene ring, forming a hydrophobic interaction with the extracellular vestibule (Fig. 2b). Structure-activity relationship studies during the development of canagliflozin showed that its inhibitory activity increases with furan replaced by thiophene in the center^25^, suggesting that this moiety should be hydrophobic. Thus, F98 and F453 are suggested to play important roles in the inhibitory action. It has been reported that SGLT1 has I98 at the position corresponding to V95 in hSGLT2 TM2, and that V95I reduced the inhibitory activity of empagliflozin in hSGLT2^19^. The distal aromatic rings also contribute to the selectivity between hSGLT1 and hSGLT2^8^.

### High concentration phlorizin fixes hSGLT2 in the inward-open state

Unexpectedly, phlorizin was found to bind to TM1, TM5, and TM8 in an inward-open structure (Fig. 3a), in contrast to the above gliflozins. This binding site is located near the Na2 site, where sodium ions bind to the outward-facing structure (Fig. 2b). The corresponding site is occluded in the inward-facing state of vSGLT and SGLT1 (Supplementary Fig. 12)^15,17^, and a PEG molecule was observed in the same location in the inward-open structure of vSGLT (Supplementary Fig. 12)^14^. The glucose moiety of phlorizin is bound to the bending site of the intracellular side of TM1, whereas the aglycon moiety, connected to the glucose moiety via an ether bond, is surrounded by the side chain of A69, S70, A73 and S74 of TM1, D201 of TM5, and R300 of TM6, extending toward the lipid membrane (Fig. 3c). The ether bond is unique to phlorizin, whereas the central aromatic ring of gliflozins connects directly to the glucose moiety, causing rigidity that prevents binding to the inward-opening structure.

The binding of phlorizin to the hSGLT2 expressing membrane fraction at up to 5000 nM did not saturate and exhibit biphasic kinetics, indicating that phlorizin has two binding sites on hSGLT2 (high- and low-affinity sites; Fig. 3d–f). Similarly, the binding of [^3^H]phlorizin exhibits biphasic binding in rat renal plasma membrane^33^. In whole-cell clamp experiments, hSGLT2 inhibitors, including phlorizin, achieve inhibition by acting on the extracellular side^34,35^; but phlorizin is the only one that has been reported to act weakly from the intracellular side at high concentrations^34^. Inhibition from the inside is more potent when there is no intracellular or extracellular Na^+^ concentration gradient^34^. Here, we added high concentration (500 μM) of phlorizin to the hSGLT2–MAP17 solution during the purification process, with no Na^+^ gradient; cryo-EM captured the binding state of phlorizin accessing from the intracellular side. We examined the binding activity and transport function of hSGLT2 alanine mutants of residues S74 or D201 involved in phlorizin binding in the inward-open structure, and of F98 or F453 involved in the inhibitor interactions in the outward-facing structure of hSGLT2. Based on FSEC, all mutants (including the WT) preserved their conformation (Supplementary Fig. 1d).

Unexpectedly, the F98A and F453A single mutants did not bind to phlorizin (Fig. 3e); F98 and F453 not only participate in the inhibitor binding in the outward-facing conformation, but also form π-π stacking interactions with each other in the inward-open conformation, which is probably important for maintaining the inward-open state (Fig. 3c). The low-affinity binding phase of phlorizin was lost in both the S74A and D201A mutants, with phlorizin binding only to the high-affinity site of the WT (Fig. 3e, f, Supplementary Data Table 3). Phloretin, the aglycon tail of phlorizin, binds to the WT as well as phlorizin but not to the S74A and D201A mutants (Fig. 3g). Therefore, S74A and D201A have lost the ability to bind phlorizin on the cytoplasmic side but can still bind it on the extracellular site. Since the S74A, F98A, and F453A mutants maintained α-MG uptake (Fig. 3h), we performed experiments to inhibit sugar uptake using phlorizin. Inhibition of α-MG uptake by phlorizin was greatly impaired in the F98A and F453A mutants, which lacked phlorizin-binding ability, but was maintained in the S74A mutant, in which the outer binding site was functional (Fig. 3i, Supplementary Data Table 4). No clear uptake of α-MG was observed in the D201A mutant (Fig. 3h). D201 corresponds to D204 in hSGLT1, which is involved in the Na3 site formation and is important for sugar uptake activity and cell trafficking; it is expected to play a similar role in hSGLT2^36^. These results support the previous suggestions that phlorizin strongly and weakly inhibits hSGLT2 from the extracellular and intracellular sides, respectively.

### Structural rearrangement from the outward to inward conformations

Given that hSGLT2 exhibits outward and inward conformations, we will consider its structural changes during sugar transport. TM1-10 is conserved in the LeuT family. hSGLT2 contains a bundle domain comprising TM1, −2, −6, and −7, a hash domain comprising TM3, −4, −8, and −9, and two gating helices, TM5 and −10. In addition, the core of hSGLT2 was surrounded by TM0, TM11–13, and MAP17 (Fig. 4).

The bundle domain of LeuT-fold transporters such as Mhp1 and vSGLT1 is reportedly fixed, while the hash domain and gating helices rotate to transport substrates via an alternating-access mechanism^12^. When the bundle domains are superimposed between the inward and outward structures, TM13 and MAP17 are also well superimposed, while the other parts of the transporter change their location and conformation (Fig. 4a, Supplementary Movie 1). MAP17 is expected to affect the active conformation of hSGLT2, because its co-expression enhances hSGLT2 activity, without altering hSGLT2 expression on the plasma membrane^18^. However, the present structure suggests that MAP17 stabilizes the bundle domain together with TM13, as a scaffolding protein in the plasma membrane (Fig. 4a).

The substrate-binding site and external vestibule are formed by TM1, −2, −6, and −10 in the outward-facing conformation (Fig. 4b, c). Y290 in TM6, N75 in TM1, and K154 in TM3 form π-cation interactions, where the corresponding interaction of vSGLT1 is thought to act as an inner gate for substrates. SGLT1/2 has a characteristic Pro-Pro motif (465, 466) in the middle of TM10, causing a bend in the α-helix (Fig. 4c, e). This causes F453 of TM10 to form a T-shaped π–π interaction with F98 of TM2 in the inward-open conformation; the external vestibule is covered by this interaction, and by L84 of TM1 (Fig. 4e). These residues are thought to act as external gates for transporters. The binding of the distal aromatic ring to the outward-facing conformation moves the F453 side chain of TM10 to the opposite side, and instead the distal aromatic ring forms a tight interaction with F98 of TM2, thus inhibiting the inward transition (Supplementary Movie 1). This is consistent with the reduced binding activity of SGLT2 inhibitors in the F98A and F453A mutants (Supplementary Fig. 1e).

The inward-open structure, which has no Na^+^ bound, is thought to mimic the structure once Na^+^ and the substrate are transported into the cytoplasmic side^14^. With the movement of the hash domain and gating helices, the interaction between K154 of TM3 and Y290 of TM6 is broken (Fig. 4c, e), and TM8 moves so that it fills the space partially occupied by TM3. Furthermore, TM8 and the intracellular part of its connecting TM9 outwardly shift significantly. The small-loop structure of IL0 stabilizes the outward-facing structure from the intracellular side, together with a cation-π interaction between R427 and Y55, which becomes lost and disordered in the inward-open conformation. In SGLT1 and vSGLT, the small helical structure IL0 in the inward-open conformation is also disordered ^14,17^, and is therefore thought to be conserved among these proteins.

Because phlorizin was bound to the low-affinity binding site in the inward-open conformation under these experimental conditions, we suggest that, in the absence of an Na^+^ concentration gradient, hSGLT2 is stabilized in the inward-open conformation, and that the Na^+^ concentration gradient may promote sodium binding to the Na2 site. Although this study does not fully elucidate the dynamics of Na^+^ binding/release and sugar uptake, it reveals that sugar uptake depends on the change from the outward-facing to the inward-open conformation, and that groove formation by the inward-open structure without Na^+^ binding is the driving force of sugar uptake. Based on molecular dynamics studies of vSGLT, sugar uptake occurs after Na^+^ is released^14^, which is consistent with our findings. After the uptake of sugar and sodium is completed in the inward-open state of the protein, the sodium concentration gradient drives the change toward the outward-open conformation, allowing the next sodium ion to be accepted, thus rotating the glucose transport cycle. Our structural findings therefore provide support for the proposed Na^+^-glucose co-transport mechanism.

In summary, we have elucidated the structures of five hSGLT2–MAP17-inhibitor complexes using cryo-EM, in addition to the sodium-binding outward-facing and inward-open structures of hSGLT2 and a two-phase mode of inhibition. This allowed us to identify the unprecedented role of Na^+^ ions in regulating transport state dynamics (Fig. 5). Most sodium-bound symporters of the LeuT family share their Na2 site and domain structures and may employ a common transport mechanism. We believe that our findings will help us to better understand the molecular mechanism of action of this transporter family and to develop new drugs that target disease-related transporters.

**Figure 5.**
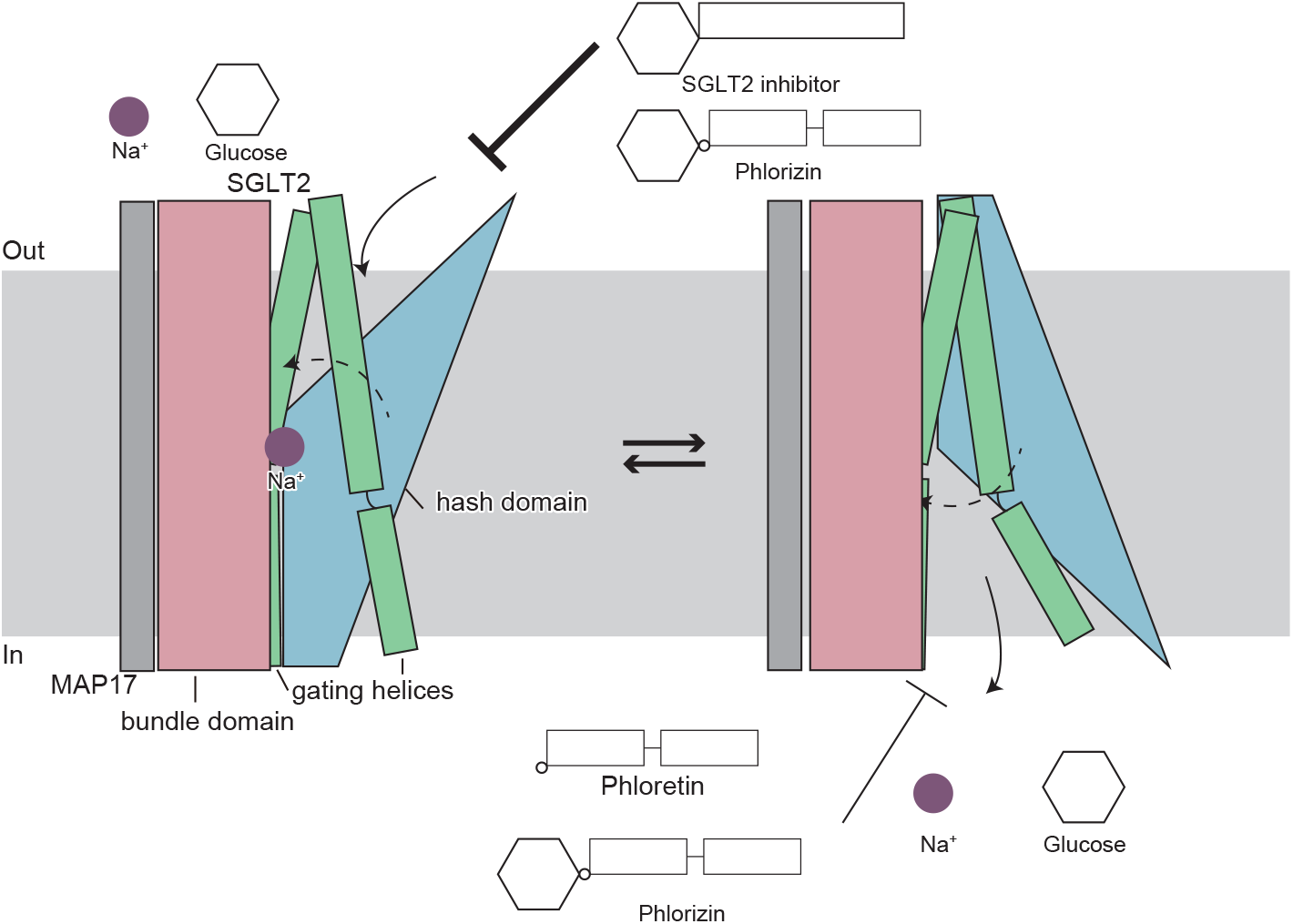
Proposed SGLT2 transport and inhibition mechanism. The bundle domain is anchored to the membrane with MAP17, and the rest of the transporter undergoes a conformational change according to an alternating-access mechanism. Upon sodium binding, the transporter opens outward to allow a substrate or inhibitors to bind. Upon substrate binding, the inner gate opens and sodium and glucose are released into the cell. After sodium and glucose are released, the transporter forms an inward-open structure and phlorizin and phloretin bind to this structure, inhibiting glucose transport.

## Methods

### Reagent and Chemicals

Canagliflozin and TA-1887 were synthesized by Mitsubishi Tanabe Pharma Corporation (Yokohama, Japan)^32,37^. Dapagliflozin and sotagliflozin were purchased from Cayman Chemical Company (Ann Arbor, MI). Phlorizin, phloretin, and α-MG were purchased from Sigma-Aldrich (St. Louis, MO).

### Expression and purification of the hSGLT2-MAP17 heterodimer

hSGLT2 (UniProt: P31639-1) cDNA and human MAP17 (UniProt: Q13113-1) cDNA were synthesized and codon-optimized for expression in human cell lines. Both cDNAs were cloned into pcDNA3.4 vector. The hSGLT2 sequence was fused with an N-terminal signal sequence from human trypsinogen 1, a His10 tag, and super-folder green fluorescent protein (sfGFP), followed by a human rhinovirus 3C protease (HRV3C protease) recognition site. Point mutations were introduced into this construct using site-directed mutagenesis.

Mammalian Expi293 cells (Thermo Fisher Scientific, Waltham, MA) were grown and maintained in Expi293 Expression Medium at 37 °C and 8% CO_2_ under humidified conditions. Cells were transiently transfected at a density of 2.0 × 10^6^ cells mL^−1^ with the plasmids and FectoPRO (Polyplus, Illkirch-Graffenstaden, France). Approximately 320 μg of the hSGLT2 plasmid and 160 μg of the MAP17 plasmid were premixed with 720 μL of FectoPRO reagent in 60 mL of Opti-MEM (Gibco) for 10–20 min before transfection. For transfection, 60 ml of the mixture was added to 0.6 L the cell culture and incubated at 37 °C in the presence of 8% CO_2_ for 72 h before collection. The cells were collected by centrifugation (800 *×g*, 10□min, 4□°C) and stored at −80 °C before use. The detergent-solubilized proteins were analyzed by fluorescence-detection size-exclusion chromatography (FSEC) using an ACQUITY UPLC BEH450 SEC 2.5 μm column (Waters, Milford, MA).

To prepare the complex sample with phlorizin, the cells were solubilized for 1 h at 4□°C in buffer (50□mM HEPES-NaOH [pH 7.5], 300□mM NaCl, 2% (w/v) N-dodecyl β-d-maltoside (DDM, Calbiochem, San Diego, CA), protease inhibitor cocktail, and 1 mM phlorizin). After ultracentrifugation (138,000 *×g*, 60□min, 4□°C), the supernatant was incubated with Affi-Gel 10 (Bio-Rad, Hercules, CA) coupled with a GFP-binding nanobody^38^ and incubated for 2□h at 4□°C. The resin was washed five times with three column volumes of wash buffer (50 mM HEPES-NaOH (pH 7.5), 300 mM NaCl, 0.05% DDM (GLYCON Biochemicals, Luckenwalde, Germany), and 1 mM phlorizin), and gently suspended overnight with HRV3C protease to cleave the His10-sfGFP tag. After HRV3C protease digestion, the flow-through was pooled, concentrated, and purified by size-exclusion chromatography on a Superose 6 Increase 10/300 GL column (GE Healthcare, Chicago, IL), and equilibrated with SEC buffer (20□mM HEPES-NaOH [pH 7.5], 150□mMNaCl, 0.03% DDM [GLYCON], and 0.5 mM phlorizin). For the complex samples with canagliflozin, TA-1887, dapagliflozin, and sotagliflozin, the same procedure was performed, but at inhibitor concentrations of 30 μM each. The peak fractions were pooled and concentrated to 6–10□mg□ml^−1^.

### Methyl α-D-glucopyranoside (α-MG) uptake in hSGLT2-transfected HEK293 cells

HEK293 cells (ECACC 85120602) were maintained in Dulbecco’s modified Eagle’s medium (Gibco) supplemented with 10% fetal bovine serum (FBS; Thermo Scientific), 2 mM l-glutamate, 100 U/mL benzylpenicillin, and 100 μg/mL streptomycin, at 37 °C in a humidified atmosphere of 5% CO_2_ in air. HEK293 cells were transiently transfected with 0.25 μg pcDNA3.4 vector containing a MAP17-coding region and 0.50 μg pcDNA3.4 vector containing an hSGLT2-coding region, using Lipofectamine 2000 (Life Technologies, Carlsbad, CA), and cultured for 48 h. The medium was removed, and the cells were washed twice, then preincubated with extracellular fluid buffer without glucose (122 mM NaCl, 25 mM NaHCO_3_, 3 mM KCl, 1.4 mM CaCl_2_, 2 mM MgSO_4_, 0.4 mM K_2_HPO_4_, 10 mM HEPES; pH 7.4) at 37 °C for 20 min. After preincubation, uptake was initiated by replacing the preincubation buffer with extracellular fluid buffer containing 500 μM α-MG in the absence or presence of inhibitors. Uptake was completed by removing the uptake buffer and washing with ice-cold buffer three times, followed by solubilization in 1 N NaOH at room temperature. The uptake of α-MG gradually increased over 60 min (Supplementary Fig. 1E) and the incubation time in the inhibition assay was 30 min.

Cell lysates were deproteinized by adding acetonitrile containing candesartan as the internal standard. The concentration of α-MG was quantified via LC-MS/MS, using the internal standard method.

Specific peaks of α-MG were observed in the lysates of mock cells and hSGLT2-expressing cells incubated with α-MG, but no specific peaks were observed in the lysates of mock cells without the addition of α-MG (Supplementary Fig. 1F). Cellular protein content was determined using a bicinchoninic acid (BCA) protein assay kit (Thermo Fisher Scientific). The uptake of α-MG was expressed as the ratio of pmol/mg protein in the cells to pmol/μL in the medium (cell-to-medium ratio; μL/mg protein).

In the inhibition study, the cell-to-medium ratio of cells transfected with the empty vector was used as the background. The specific α-MG uptake was calculated by subtracting the background from the total cell-to-medium ratio and normalized to the uptake achieved without the inhibitor. The three replicates were obtained from the same samples. IC50 was calculated via nonlinear regression using GraphPad Prism 8.4.3.

### SGLT2 inhibitor-binding assay by affinity selection-mass spectrometry

To examine the inhibition of binding to the crude membrane, mammalian Expi293 cells were co-transfected with hMAP17 and wild-type hSGLT2 or its mutants, as described above. The cells were collected and disrupted by sonication in a hypotonic buffer (50□mM HEPES-NaOH [pH 7.5], 10□mM KCl, and protease inhibitor cocktail) or Na^+^-free hypotonic buffer (50□mM Tris-HCl [pH 7.5], 10□mM KCl, and protease inhibitor cocktail). Cell debris were removed by centrifugation (2,000□*×g*, 5□min, 4□°C). The membrane fraction was collected by ultracentrifugation (112,000 *×g*, 30 min, 4□°C) and stored at −80 °C before use. The crude membrane (250 μg per sample) was incubated with SGLT2 inhibitor in an assay buffer (100 mM NaCl, 10 mM HEPES/Tris, pH 7.4) or Na^+^-free assay buffer (100 mM choline chloride, 10 mM HEPES/Tris, pH 7.4) at room temperature for 2 h. Reactions were terminated by filtration through a GF/C filter plate (Corning Inc., Corning, NY) presoaked in assay buffer containing 0.1% BSA. The sample in the filter plate was washed three times with the assay buffer and eluted with acetonitrile: water (80:20, v/v). The extract solution from the filter plate was diluted with water containing candesartan as an internal standard, and the concentration of SGLT2 inhibitors was quantified using LC-MS/MS.

Non-specific binding was determined by the membrane fraction of the non-transfected-Expi293 cells, and specific binding was calculated by subtracting the non-specific binding from the binding of hSGLT2-expressing cells. Specific binding was normalized to hSGLT2 protein expression levels, measured by FSEC. The equilibrium dissociation constant (Kd) and maximum number of binding sites (B_max_) were calculated via nonlinear regression in GraphPad Prism 8.4.3. The specific binding of the hSGLT2 mutants was normalized to that of wild-type hSGLT2.

### Quantification of SGLT2 substrate and inhibitors via LC-MS/MS

The concentration of the extract solution from the filter plate and cell lysate was quantified using tandem mass spectrometry (QTRAP6500 System (SCIEX, Framingham, MA) coupled with an ACQUITY UPLC system (Waters) using the internal standard method. Mobile phases A and B used 10 mM of ammonium bicarbonate and acetonitrile, respectively. Chromatographic separation was performed on an ACQUITY UPLC BEH C18 column (2.1 mm × 100 mm, 1.7 μm; Waters) at 50 °C, with the following gradient of mobile phase B: 1% (at 0.00 to 0.50 min), 1% to 95% (0.50 to 2.00 min), 95% (2.00 to 2.50 min), and 1% (2.51 to 3.0 min); the flow rate was 0.4 mL/min. Mass spectrometric detection was performed by multiple reaction monitoring in the electrospray-ionization negative-ion mode, using m/z 443.1 / 364.9 for canagliflozin; 425.9 / 264.1 for TA-1887; 407.0 / 328.8 for dapagliflozin; 423.0 / 387.0 for sotagliflozin; 435.0 / 273.0 for phlorizin; 273.0 / 148.9 for phloretin; and 192.9 / 100.9 for α-MG.

### Electron microscopy sample preparation

The purified protein solution of hSGLT2–MAP17 was mixed with the inhibitor solutions except for phlorizin, with final concentrations of 0.5 mM dapagliflozin, TA-1887, sotagliflozin, or canagliflozin. After incubation for 1 h on ice, the grids were glow-discharged in low-pressure air with a 10□mA current in a PIB-10 (Vacuum Device, Mito, Japan). The protein solutions containing 0.5 mM of the inhibitors were applied to a freshly glow-discharged Quantifoil Holey Carbon Grid (R1.2/1.3, Cu/Rh, 300 mesh)(SPT Labtech, Melbourne Hertfordshire, UK) using a Vitrobot Mark IV (Thermo Fisher Scientific) at 4 °C, with a blotting time of 4–6 s under 99% humidity conditions; the grids were then plunge-frozen in liquid ethane.

### Electron microscopy data collection and processing

The grids containing phlorizin, TA-1887, dapagliflozin, and sotagliflozin were transferred to a Titan Krios G3i (Thermo Fisher Scientific) running at 300 kV and equipped with a Gatan Quantum-LS Energy Filter (GIF) and a Gatan K3 Summit direct electron detector in the correlated double sampling mode. Imaging was performed at a nominal magnification of 105,000×, corresponding to a calibrated pixel size of 0.83 Å/px (The University of Tokyo, Japan). Each movie was dose fractionated to 64 frames at a dose rate of 6.2–9.0 e^−^/px/s at the detector, resulting in a total accumulated exposure of 64 e^−^/Å^2^ of the specimen. The data were automatically acquired using the image-shift method in SerialEM software^39^, with a defocus range of −0.8 to −1.6 μm.

The grid with canagliflozin was transferred to a Titan Krios G4 device (Thermo Fisher Scientific) running at 300 kV and equipped with a Gatan Quantum-LS Energy Filter (GIF) and a Gatan K3 Summit direct electron detector in correlated double sampling mode. Imaging was performed at a nominal magnification of 215,000×, corresponding to a calibrated pixel size of 0.4 Å/px (The University of Tokyo, Japan). Each movie was recorded for 1.4 seconds and subdivided into 64 frames. Electron flux was set to 7.5 e^−^/px/s at the detector, resulting in an accumulated exposure of 64 e^−^/Å^2^ of the specimen. The data were automatically acquired via the image-shift method using EPU software (Thermo Fisher Scientific), with a defocus range of −0.6 to −1.6 μm. The total number of images is described in Supplementary Data Table 1.

For all datasets, the dose-fractionated movies were subjected to beam-induced motion correction using RELION^23^, and the contrast transfer function (CTF) parameters were estimated using CTFFIND4^40^.

For the canagliflozin-bound state dataset, 2,364,108 particles were initially selected from 19,943 micrographs, using the topaz-picking function in RELION-4.0^24^. Particles were extracted by downsampling to a pixel size of 3.2 Å/px. These particles were subjected to several rounds of 2D and 3D classification. The best class contained 221,701 particles, which were then re-extracted with a pixel size of 1.60 Å/px and subjected to 3D refinement. The second 3D classification resulted in three map classes. The best class from the 3D classification contained 179,761 particles, which were subjected to 3D refinement. The particles were subsequently subjected to micelle subtraction and non-aligned 3D classification using a mask (without micelles), resulting in three map classes. The best class, containing 65,919 particles, was subjected to 3D refinement, reversion to the original particles, and 3D refinement. The particle set was resized to 1.00 Å/px, and subjected to Bayesian polishing, 3D refinement, and per-particle CTF refinement before the final 3D refinement and post-processing, yielding a map with a global resolution of 3.1 Å, according to the FSC = 0.143 criterion. Finally, local resolution was estimated using RELION-4. The processing strategy is illustrated in Supplementary Fig. 3.

For the dapagliflozin-bound-state dataset, 3,692,950 particles were initially selected from 4,841 micrographs, using the topaz-picking function in RELION-4.0. Particles were extracted by downsampling to a pixel size of 3.32 Å/px. These particles were subjected to several rounds of 2D and 3D classification. The best class contained 569,516 particles, which were then re-extracted at a pixel size of 1.30 Å/px and subjected to 3D refinement. Non-aligned 3D classification using a soft mask covering the proteins and micelles resulted in four map classes. The best class from the 3D classification contained 197,695 particles, which were subjected to 3D refinement, per-particle CTF refinement, and 3D refinement. The resulting 3D model and particle set were resized to 1.11 Å/px and subjected to Bayesian polishing, 3D refinement, and per-particle CTF refinement. Final 3D refinement and post-processing yielded maps with global resolutions of 2.8 Å, according to the FSC = 0.143 criterion. Finally, the local resolution was estimated using RELION-3. The processing strategy is illustrated in Supplementary Fig. 4.

For the TA-1887-bound state dataset, 3,395,470 particles were initially selected from 4,383 micrographs, using the topaz-picking function in RELION-4. Particles were extracted by downsampling to a pixel size of 3.32 Å/px. These particles were subjected to several rounds of 2D and 3D classification. The best class contained 274,477 particles, which were then re-extracted with a pixel size of 1.30 Å/px and subjected to 3D refinement. Non-aligned 3D classification using a soft mask covering the proteins and micelles resulted in three map classes. The best class from the 3D classification contained 103,853 particles, which were subjected to 3D refinement, per-particle CTF refinement, and 3D refinement. The resulting 3D model and particle set were resized to 1.11 Å/px and subjected to Bayesian polishing, 3D refinement, and per-particle CTF refinement. Final 3D refinement and post-processing yielded maps with global resolutions of 2.9 Å, according to the FSC = 0.143 criterion. Local resolution was estimated using RELION-4. The processing strategy is illustrated in Supplementary Fig. 5.

For the sotagliflozin-bound state dataset, 5,242,427 particles were initially selected from 5,499 micrographs using the topaz-picking function in RELION-4. Particles were extracted by downsampling to a pixel size of 3.32 Å/px. These particles were subjected to several rounds of 2D and 3D classifications. The best class contained 823,369 particles, which were then re-extracted with a pixel size of 1.30 Å/px and subjected to 3D refinement. Non-aligned 3D classification using a soft mask covering the proteins and micelles resulted in four classes of maps. The two good classes from the 3D classification contained 227,811 particles, which were subjected to 3D refinement. The resulting 3D model and particle set were resized to 1.11 Å/px and subjected to Bayesian polishing, 3D refinement, and further non-aligned 3D classification using a soft mask covering the proteins and micelles. The best class from the 3D classification contained 72,773 particles, which were subjected to 3D refinement, per-particle CTF refinement, 3D refinement, Bayesian polishing, 3D refinement, and per-particle CTF refinement. The final 3D refinement and post-processing yielded maps with global resolutions of 3.1 Å, according to the FSC = 0.143 criterion. Finally, the local resolution was estimated using RELION. The processing strategy is illustrated in Supplementary Fig. 6.

For the phlorizin-bound state dataset, 3,013,029 particles were initially selected from 3,159 micrographs using the Laplacian-of-Gaussian picking function in RELION-3.1^23^ and were used to generate two-dimensional (2D) models for reference-based particle picking. Particles were extracted by downsampling to a pixel size of 3.32 Å/px. These particles were subjected to several rounds of 2D and 3D classification. The best class contained 324,355 particles, which were then re-extracted with a pixel size of 1.66 Å/px and subjected to 3D refinement. The particles were subsequently subjected to micelle subtraction and non-aligned 3D classification using a mask (excluding the micelles), resulting in three map classes. The best class contained 76,485 particles, which were then subjected to 3D refinement and reversion to the original particles. The particle set was resized to 1.30 Å/px, and subjected to Bayesian polishing, 3D refinement, and per-particle CTF refinement before the final 3D refinement and post-processing, yielding a map with a global resolution of 3.3 Å according to the FSC = 0.143 criterion. Finally, the local resolution was estimated using RELION-3. The processing strategy is illustrated in Supplementary Fig. 8.

### Model building and validation

The models of the phlorizin-bound inward state of hSGLT2–MAP17 were manually built, de novo, using the cryo-EM density map tool in COOT, facilitated by an hSGLT2-homology model generated using Alphafold2^41^. After manual adjustment, the models were subjected to structural refinement via the Servalcat pipeline in REFMAC5^42^ and manual real-space refinement in COOT. The models of the dapagliflozin-, TA-1887-, sotagliflozin-, and canagliflozin-bound outward states were built using the Alphafold2-derived hSGLT2-homology model as the starting model. The 3D reconstruction and model refinement statistics are summarized in Supplementary Data Table 1. All molecular graphics figures were prepared using CueMol (http://www.cuemol.org) and UCSF Chimera^43^.

## Acknowledgements

We thank Y. Lee and T. Nishizawa for technical advice on sample preparation; K. Yamashita for technical advice on refinement using Servalcat; T. Saijo, T. Takahashi and Y. Kamikozawa for technical advice on functional analyses; and M. Shiotani, C. Kuriyama, and Y. Yamamoto for fruitful discussions about the inhibitory mechanism. We thank the scientific staff of the cryo-EM facility of the University of Tokyo, and especially Y. Kise, Y. Sakamaki, T. Kusakizako, H. Yanagisawa, A. Tsutsumi, M. Kikkawa, and R. Danev.

This work was not supported by external funding.

## Author contributions

M. H. designed the entire study. M.H. performed the cryo-EM analyses, with sample preparation assistance from T.K.; M.H. and T.A. designed and performed the functional analyses, with sample preparation assistance from M.H. and T.K. M.H. performed model building and model refinement, with assistance from H.K. and K.M. M.H., T.A., N. T., I.M., and O.N. wrote and edited the manuscript, with help from all other authors. M.H., T.A., I.M., and O.N. supervised the study.

## Materials and correspondence

Masahiro Hiraizumi, Ikuko Miyaguchi, and Osamu Nureki

## Data and materials availability

Cryo-EM density maps were deposited in the Electron Microscopy Data Bank under accession codes EMD-34673 (canagliflozin-bound state), EMD-34705 (dapagliflozin-bound state), EMD-34610 (TA-1887-bound state), EMD-34737 (sotagliflozin-bound state), and EMD-34823 (phlorizin-bound state). Atomic coordinates have been deposited in the Protein Data Bank under IDs 8HDH (canagliflozin-bound state), 8HEZ (dapagliflozin-bound state), 8HB0 (TA-1887-bound state), 8HG7 (sotagliflozin-bound state), and 8HIN (phlorizin-bound state).

The datasets generated during and/or analysed during the current study are available from the corresponding author on reasonable request.

## Figure legends

Supplementary Figure 1. Biochemical characterization of the hSGLT2–MAP17 complex.

(a) Representative size-exclusion chromatography profile of hSGLT2–MAP17. (b) SDS-PAGE analysis of the hSGLT2–MAP17 peak fractions via size-exclusion chromatography (SEC) purification. (c) SGLT2 inhibitors (25 nM) binding to the crude membrane expressing hSGLT2 and MAP17. (d) FSEC profiles for various mutations of sfGFP-tagged hSGLT2 with MAP17. The arrows indicate the elution positions of the hSGLT2–MAP17 heterodimer and the free GFP (GFP). (e) Canagliflozin binding to the crude membrane expressing wild-type hSGLT2 and mutants. Crude membranes were incubated with 30 nM canagliflozin and binding was measured by LC-MS/MS. Data are shown as mean ± SEM (n = 3). (f) Time-course of hSGLT2-mediated α-MG uptake. α-MG uptake (500 μM) by hSGLT2-expressing cells and mock cells was examined. Each point represents mean ± SEM (n = 3). (g) Chromatograms of α-MG in the lysates of untreated mock cells (No treatment), mock cells incubated with α-MG (Mock), and hSGLT2- and MAP17-expressing cells incubated with α-MG (hSGLT2).

Supplementary Figure 2. Sequence alignment of SGLT.

Sequence alignment of hSGLT2 (UniProt: P31639), mSGLT2 (UniProt: Q92317), hSGLT1 (UniProt: P13866), *Vibrio parahaemolyticus* SGLT (UniProt: P96169), and *Proteus mirabilis* HI4320 sialic acid symporter (UniProt: B4EZY7), performed using Clustal Omega. Conserved transmembrane helices of hSGLT2 are indicated above the sequences. The similarly conserved residues are indicated by red letters. The residues at the conserved Na2 and Na3 sites of SGLT are highlighted with pink circles above or white circles below the alignment, respectively.

Supplementary Figure 3. Data processing of the canagliflozin-bound state.

(a) Representative cryo-EM image of the hSGLT2–MAP17 complex in the presence of canagliflozin. (b) Data processing workflow of single-particle image-processing and local-resolution analysis. Particles were separated into three groups via non-aligned 3D classification, with the mask (without micelles) shown in yellow. (c) Cross-validation FSC curves for map-to-model fitting. (d) Angular distributions of the final reconstruction.

Supplementary Figure 4. Data processing of the dapagliflozin-bound state.

(a) Representative cryo-EM image of the hSGLT2–MAP17 complex in the presence of dapagliflozin. (b) Data processing workflow of single-particle image-processing and local-resolution analysis. Particles were separated into four groups by non-aligned 3D classification, with the mask (without micelles) shown in transparent white. (c) Cross-validation FSC curves for map-to-model fitting. (d) Angular distributions of the final reconstruction.

Supplementary Figure 5. Data processing of the TA-1887-bound state.

(a) Representative cryo-EM image of the hSGLT2–MAP17 complex in the presence of TA-1887. (b) Data processing workflow of single-particle image-processing and local-resolution analysis. Particles were separated into three groups via non-align 3D classification, with the mask covering the proteins and micelle shown in transparent white. (c) Cross-validation FSC curves for map-to-model fitting. (d) Angular distributions of the final reconstruction.

Supplementary Figure 6. Data processing of the sotagliflozin-bound state.

(a) Representative cryo-EM image of the hSGLT2–MAP17 complex in the presence of sotagliflozin. (b) Data processing workflow of single-particle image-processing and local-resolution analysis. Particles were separated into three groups via two rounds of non-aligned 3D classification, with the mask covering the proteins and micelles shown in transparent white. (c) Cross-validation FSC curves for map-to-model fitting. (d) Angular distributions of the final reconstruction.

Supplementary Figure 7. The outward-opening model of hSGLT2–MAP17 in the density maps.

The cryo-EM density and atomic model of each segment of the outward-opening model hSGLT2–MAP17, inhibitors, and glycosylation sites, contoured to 3.0 σ, 4.0 σ, and 2.0 σ, respectively. Red spheres: water molecules around the inhibitors.

Supplementary Figure 8. Data processing of the phlorizin-bound state.

(a) Representative cryo-EM image of the hSGLT2–MAP17 complex in the presence of phlorizin. (b) Data processing workflow of single-particle image-processing and local-resolution analysis. Particles were separated into three groups by non-aligned 3D classification, with the mask covering the proteins (without micelles) shown in yellow. (c) Cross-validation FSC curves for map-to-model fitting. (d) Angular distributions of the final reconstruction.

Supplementary Figure 9. Inward-open model of hSGLT2–MAP17 in the density maps. The cryo-EM density and atomic models of each segment of the inward-open model of hSGLT2–MAP17, phlorizin, and glycosylation sites, contoured to 2.9 σ, 2.7 σ, and 2.7 σ, respectively.

Supplementary Figure 10. MAP17 and SGLT2 interaction site.

The density of the lipid molecule (orange) is observed between MAP17 and SGLT2.

Supplementary Figure 11. Sodium ion-binding sites of SGLT2

(a) The sodium ion-binding Na2 sites of dapagliflozin, canagliflozin, TA-1887, and sotagliflozin are shown. (b) Sites in SGLT2 corresponding to Na3 sites where the other sodium ion binds in SGLT1. No electron density corresponding to a sodium ion can be observed. (c, d) The sodium ion-binding Na2 sites, and the sites corresponding to Na3 sites of the phlorizin-bound inward-open conformation. The phlorizin binding site is near the Na2 and Na3 sites.

Supplementary Figure 12. Comparison of the inward conformation of other SGLT structures.

Structural comparison of sites corresponding to the intracellular phlorizin-binding sites of SGLT2. From left to right, inward-occluded conformation of hSGLT1, inward-occluded conformation of vSGLT, and inward-open conformation of vSGLT.

The structures are viewed from the membrane side.

Supplementary Data Table 1.

Data collection, processing, model refinement, and validation. Clash scores, rotamer outliers, and Ramachandran plots were calculated using the Servalcat pipeline.

Supplementary Data Table 2.

Kinetic parameters of canagliflozin binding in the membrane fraction of hSGLT2-expressing and MAP17-expressing cells, in the presence or absence of Na^+^.

Supplementary Data Table 3.

Kinetic parameters of phlorizin and phloretin binding in the membrane fraction of wild-type cells or cells expressing mutated hSGLT2 and MAP17.

Supplementary Data Table 4.

IC50 values of phlorizin in α-MG uptake by wild-type and mutant hSGLT2.

Supplementary Movie 1.

Movement of SGLT2 during sugar uptake, as predicted from the cryo-EM structures. The bundle domain (TM1, −2, −6, −7; red) functions as the axis together with MAP17 (gray). TM13 (light orange), the hash domains (TM3, −4, −8, −9; blue), and the gate helices TM5 and TM10 (green) and TM0, TM11, and −12 (light orange) move significantly, transporting the sugar to the intracellular side.

