## Supplemental for "Structural insights into the mechanism of the human SGLT2–MAP17 glucose transporter"

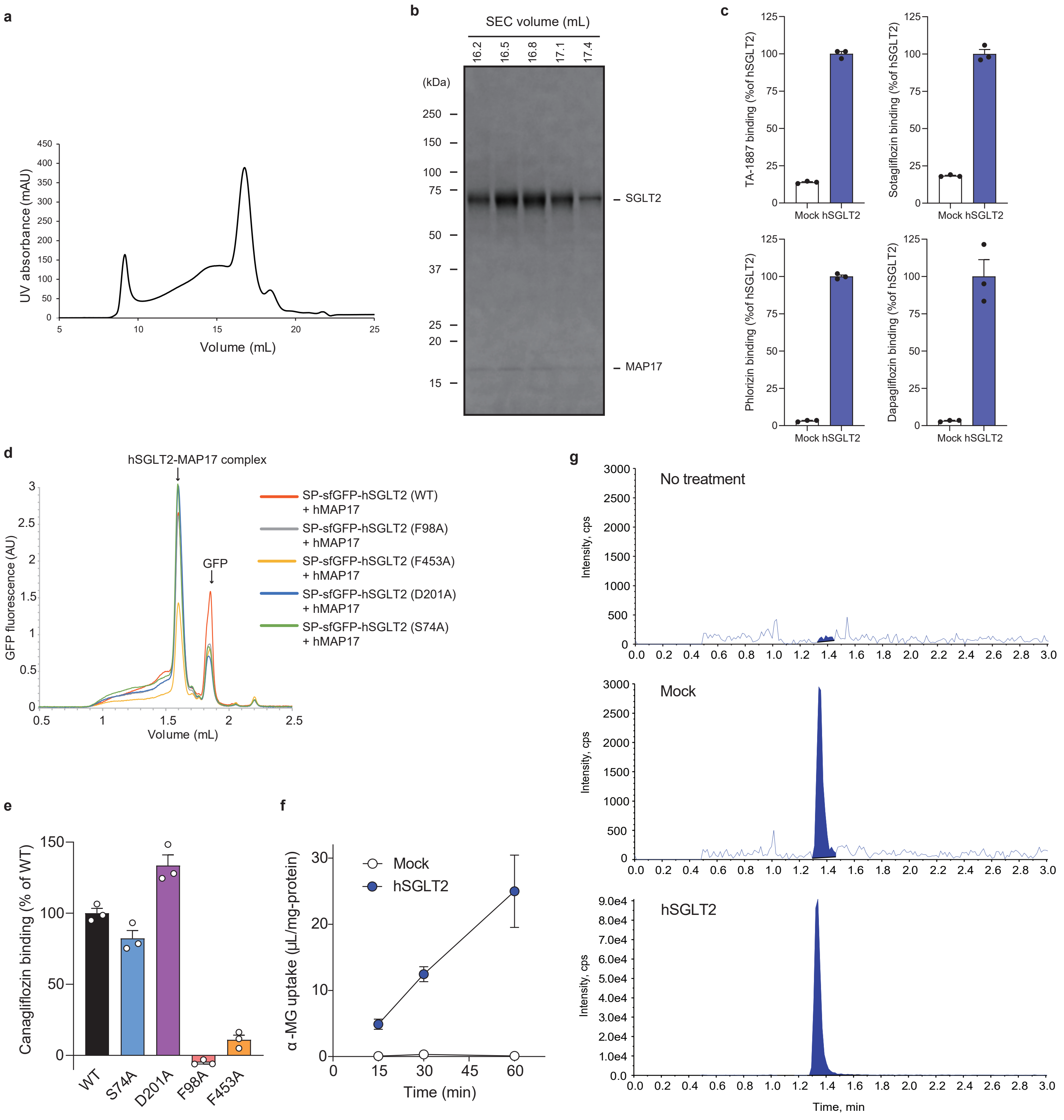


**Supplementary Figure 1. Biochemical characterization of the hSGLT2–MAP17 complex.**
(a) Representative size-exclusion chromatography profile of hSGLT2–MAP17. (b) SDS-PAGE analysis of the hSGLT2–MAP17 peak fractions via size-exclusion chromatography (SEC) purification. (c) SGLT2 inhibitors (25 nM) binding to the crude membrane expressing hSGLT2 and MAP17 (n = 3, technical replicates). (d) FSEC profiles for various mutations of sfGFP-tagged hSGLT2 with MAP17. The arrows indicate the elution positions of the hSGLT2–MAP17 heterodimer and the free GFP (GFP). (e) Canagliflozin binding to the crude membrane expressing wild-type hSGLT2 and mutants. Crude membranes were incubated with 30 nM canagliflozin and binding was measured by LC-MS/MS. Data are shown as mean ± SEM (n = 3, technical replicates). (f) Time-course of hSGLT2-mediated α-MG uptake. α-MG uptake (500 μM) by hSGLT2-expressing cells and mock cells was examined. Each point represents mean ± SEM (n = 4, biological replicates). (g) Chromatograms of α-MG in the lysates of untreated mock cells (No treatment), mock cells incubated with α-MG (Mock), and hSGLT2- and MAP17-expressing cells incubated with α-MG (hSGLT2).

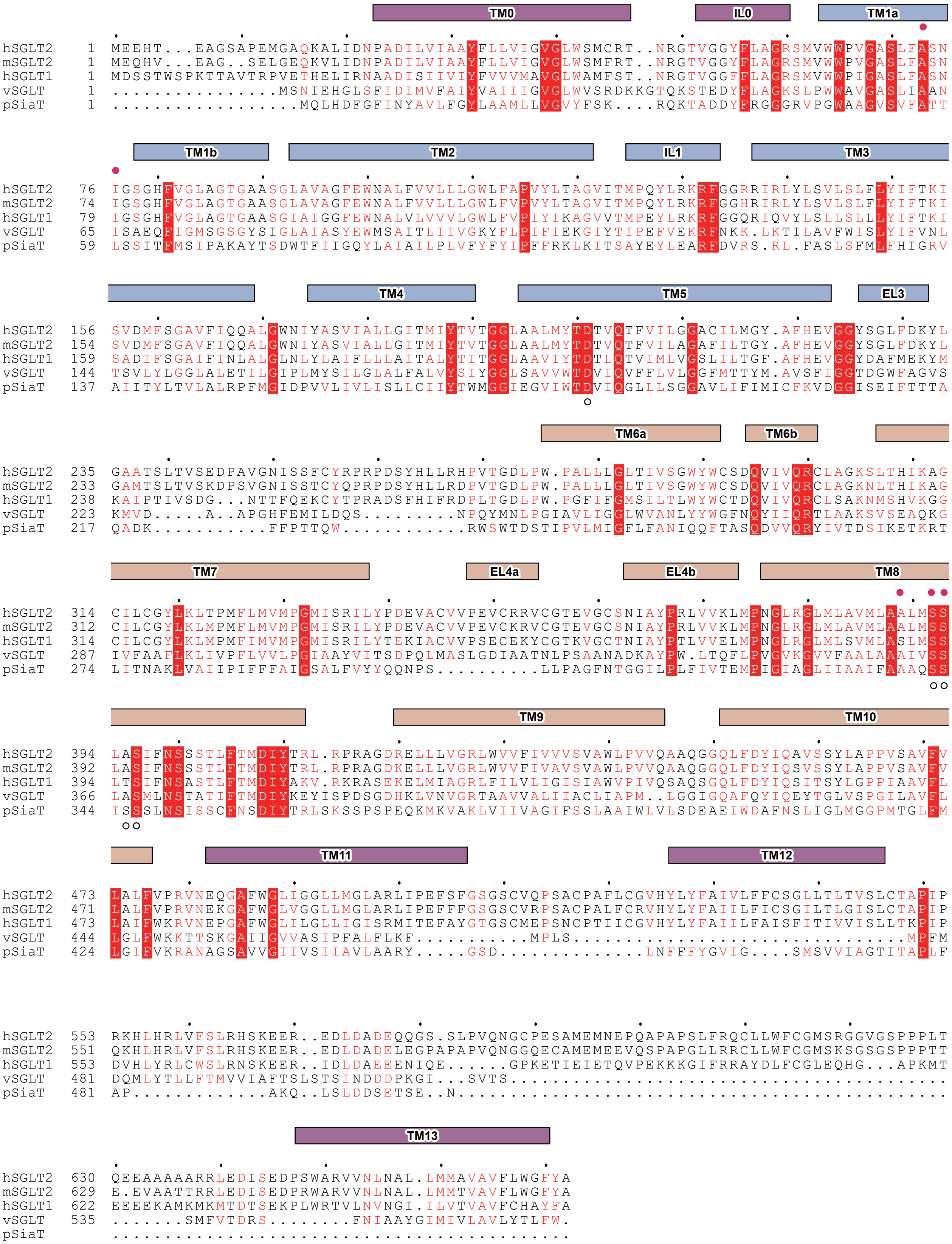


**Supplementary Figure 2. Sequence alignment of SGLT.**

Sequence alignment of hSGLT2 (UniProt: P31639), mSGLT2 (UniProt: Q92317), hSGLT1 (UniProt: P13866), Vibrio parahaemolyticus SGLT (UniProt: P96169), and Proteus mirabilis HI4320 sialic acid symporter (UniProt: B4EZY7), performed using Clustal Omega. Conserved transmembrane helices of hSGLT2 are indicated above the sequences. The similarly conserved residues are indicated by red letters. The residues at the conserved Na2 and Na3 sites of SGLT are highlighted with pink circles above or white circles below the alignment, respectively.

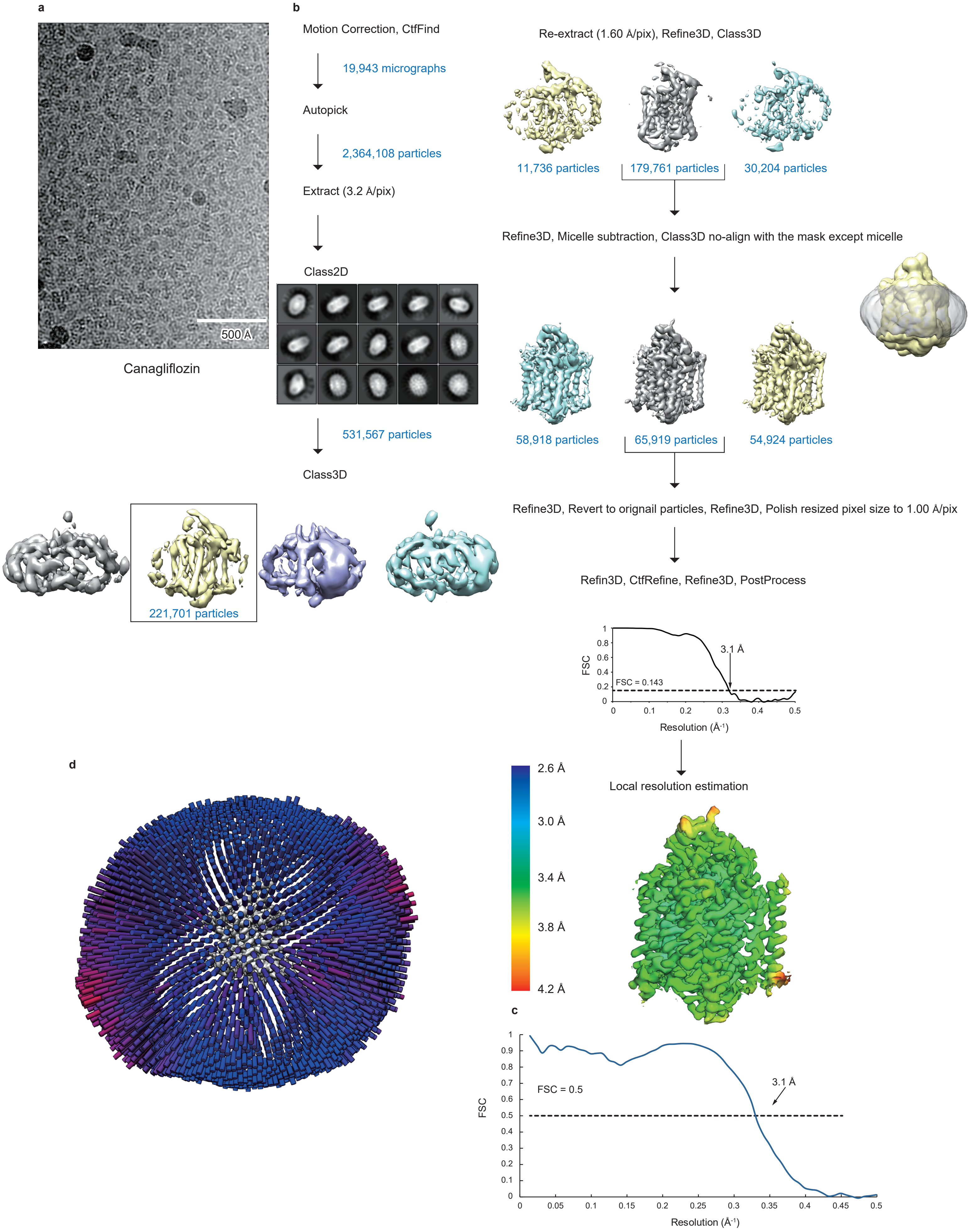


**Supplementary Figure 3. Data processing of the canagliflozin-bound state.**

(a) Representative cryo-EM image of the hSGLT2–MAP17 complex in the presence of canagliflozin. (b) Data processing workflow of single-particle image-processing and local-resolution analysis. Particles were separated into three groups via non-aligned 3D classification, with the mask (without micelles) shown in yellow. (c) Cross-validation FSC curves for map-to-model fitting. (d) Angular distributions of the final reconstruction.

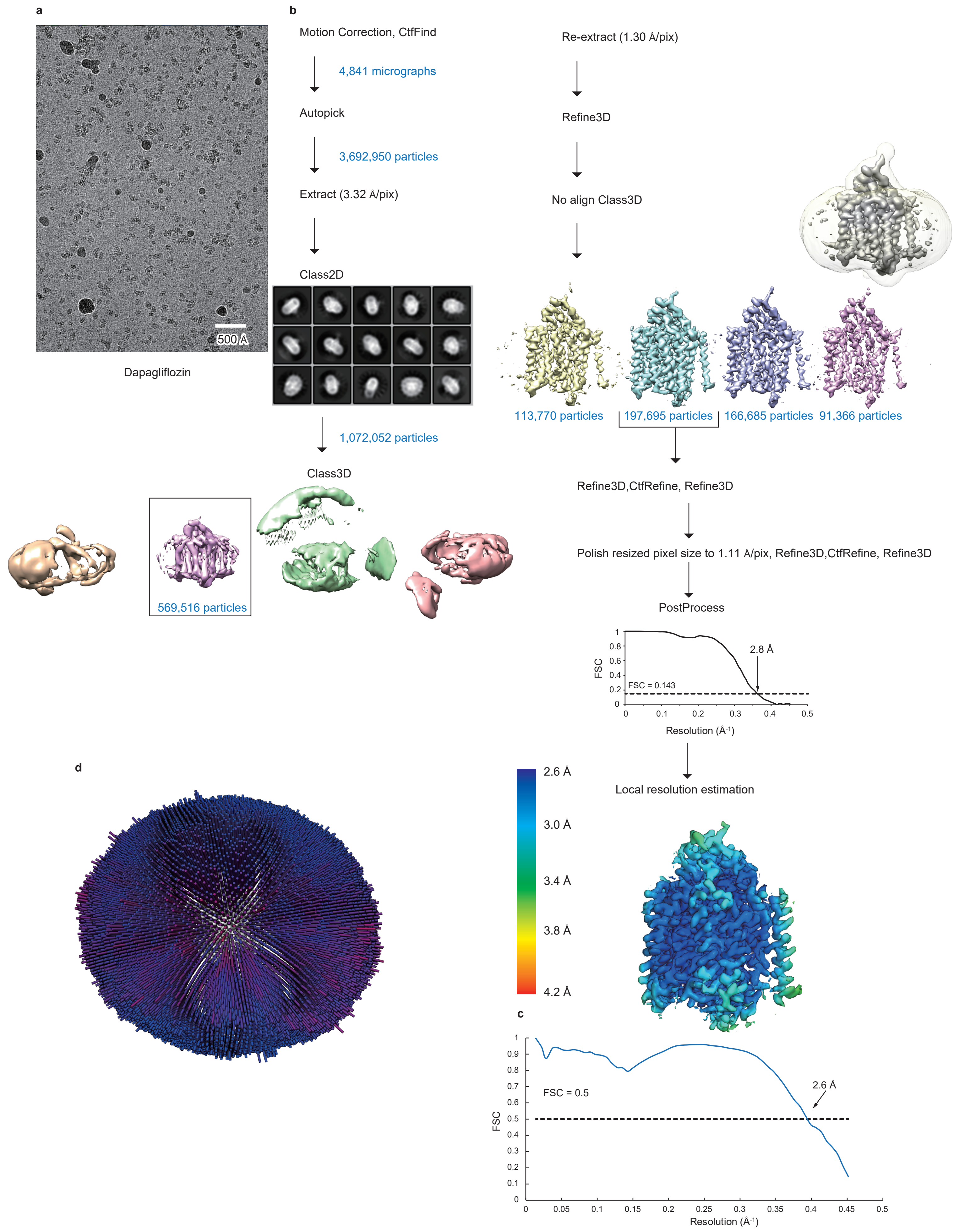


**Supplementary Figure 4. Data processing of the dapagliflozin-bound state.**

(a) Representative cryo-EM image of the hSGLT2–MAP17 complex in the presence of dapagliflozin. (b) Data processing workflow of single-particle image-processing and local-resolution analysis. Particles were separated into four groups by non-aligned 3D classification, with the mask (without micelles) shown in transparent white. (c) Cross-validation FSC curves for map-to-model fitting. (d) Angular distributions of the final reconstruction.

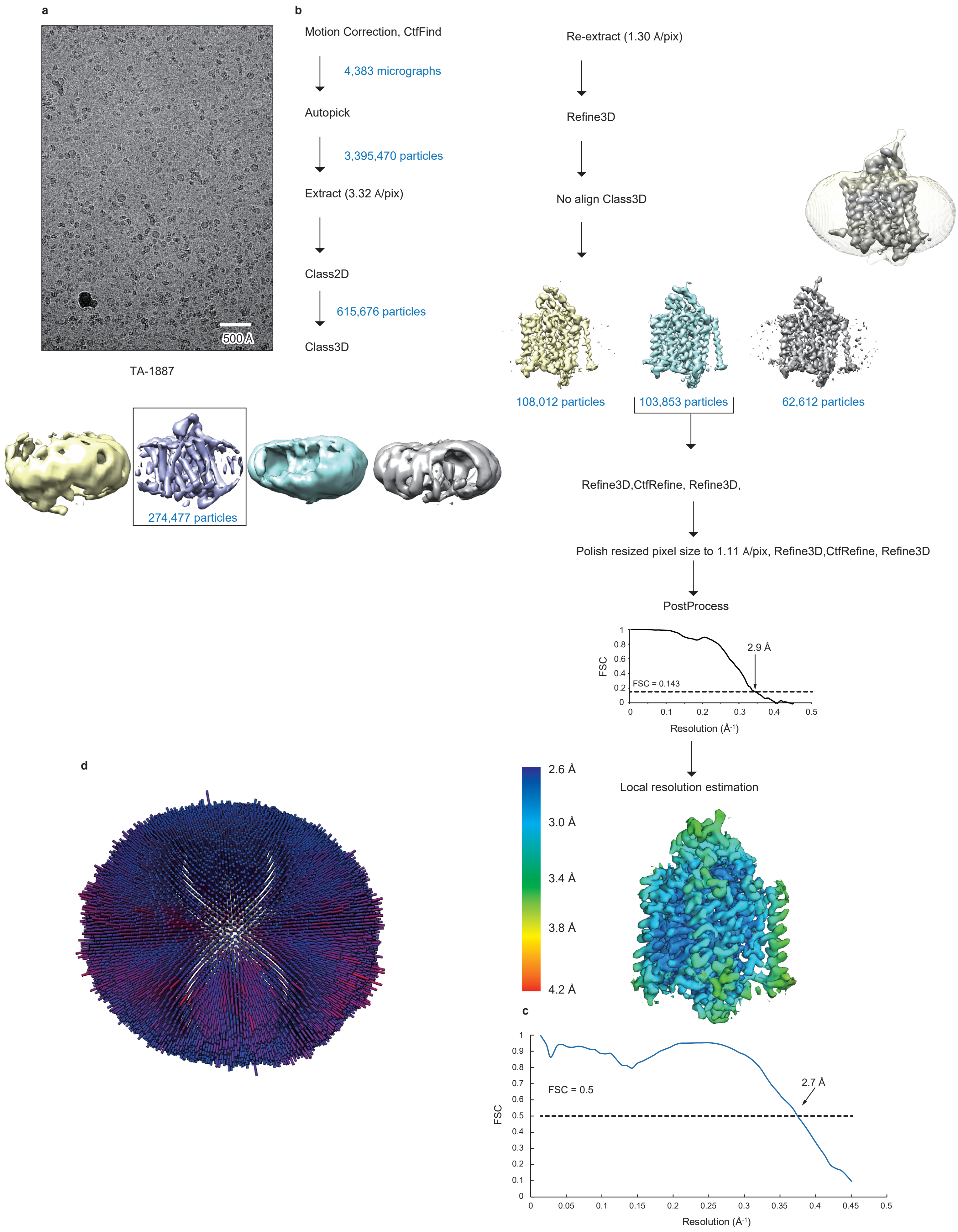


**Supplementary Figure 5. Data processing of the TA-1887-bound state.**

(a) Representative cryo-EM image of the hSGLT2–MAP17 complex in the presence of TA-1887. (b) Data processing workflow of single-particle image-processing and local-resolution analysis. Particles were separated into three groups via non-align 3D classification, with the mask covering the proteins and micelle shown in transparent white. (c) Cross-validation FSC curves for map-to-model fitting. (d) Angular distributions of the final reconstruction.

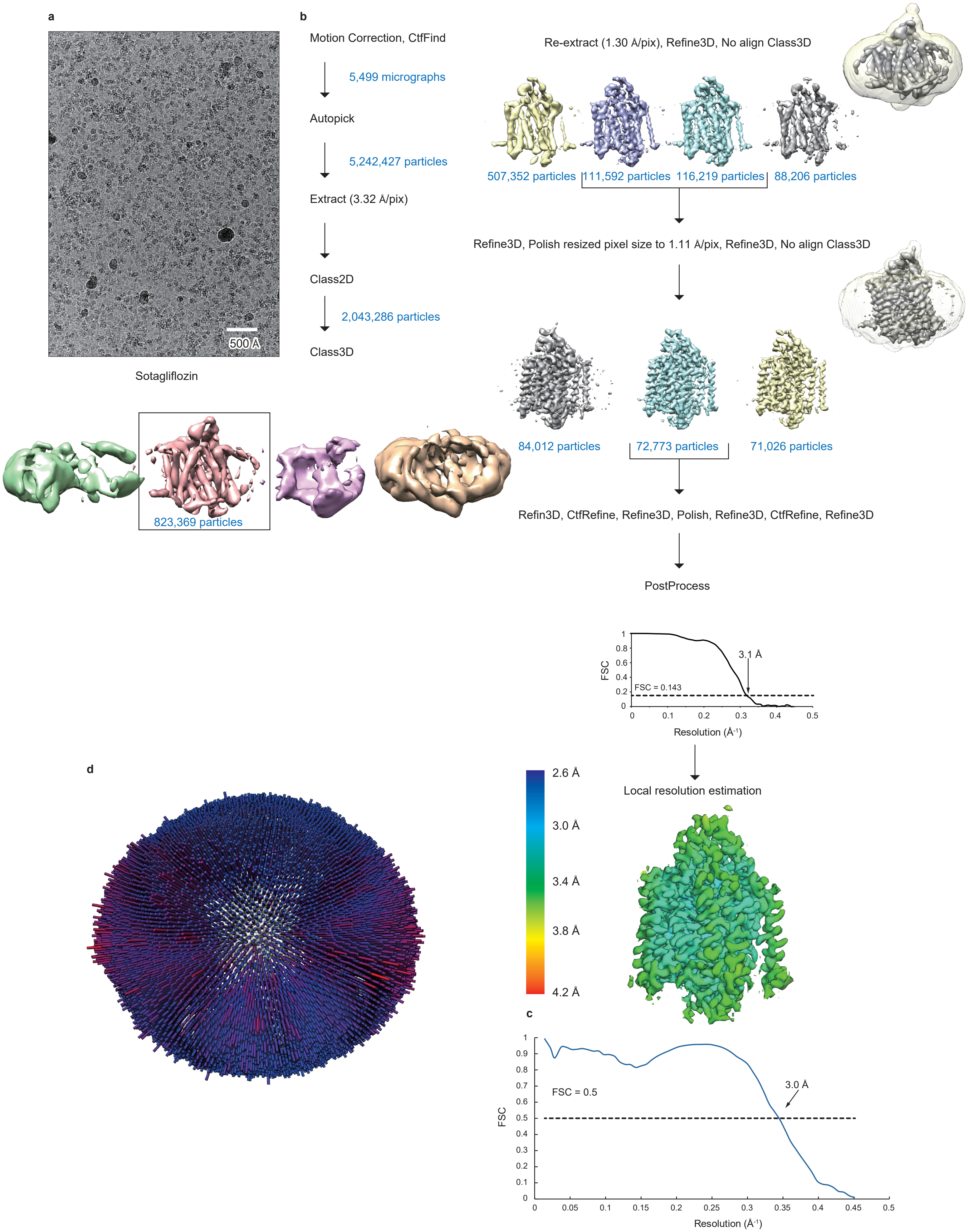


**Supplementary Figure 6. Data processing of the sotagliflozin-bound state.**

(a) Representative cryo-EM image of the hSGLT2–MAP17 complex in the presence of sotagliflozin. (b) Data processing workflow of single-particle image-processing and local-resolution analysis. Particles were separated into three groups via two rounds of non-aligned 3D classification, with the mask covering the proteins and micelles shown in transparent white. (c) Cross-validation FSC curves for map-to-model fitting. (d) Angular distributions of the final reconstruction.

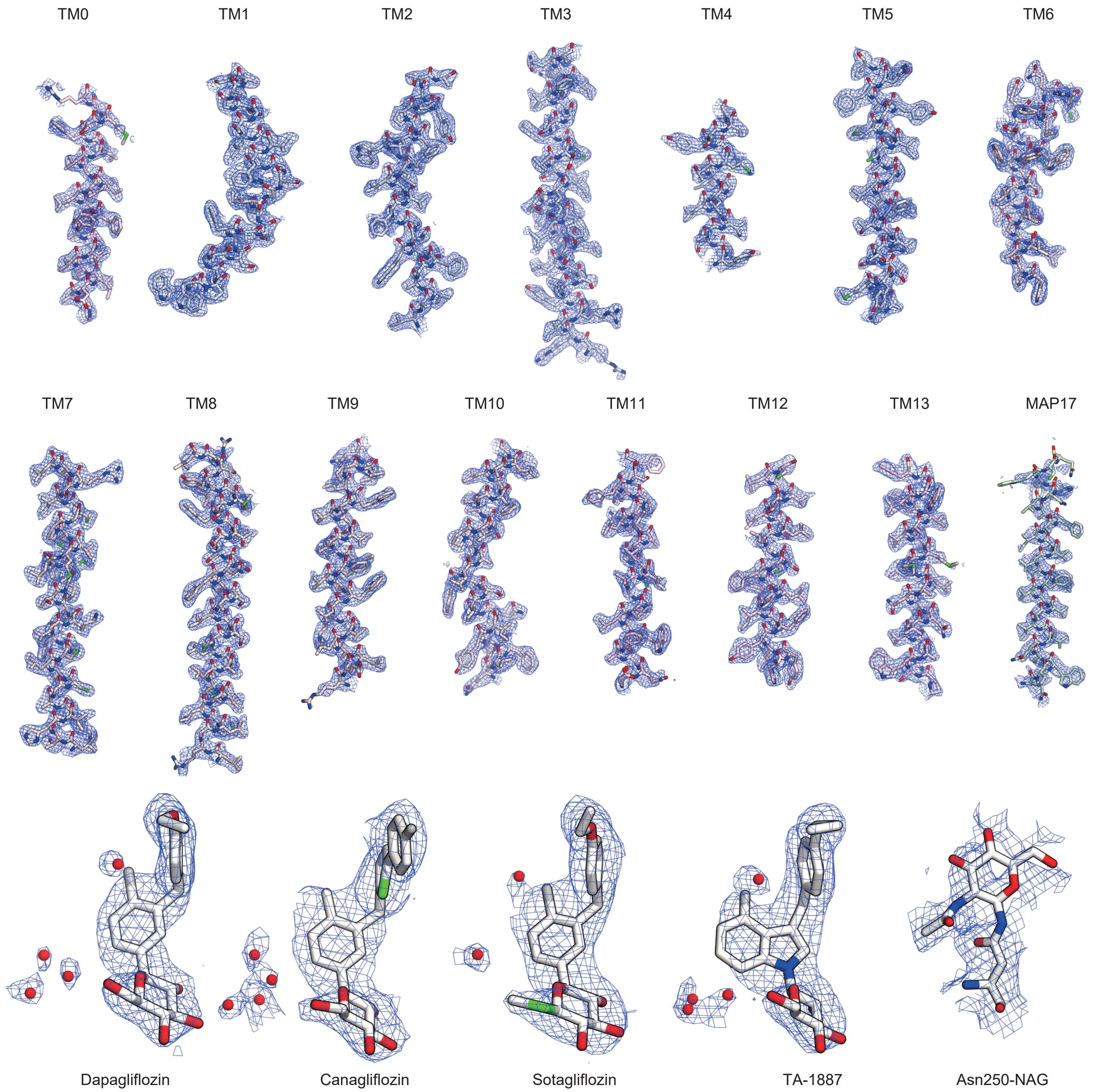


**Supplementary Figure 7. The outward-opening model of hSGLT2–MAP17 in the density maps.**
The cryo-EM density and atomic model of each segment of the outward-opening model hSGLT2–MAP17, inhibitors, and glycosylation sites, contoured to 3.0 σ, 4.0 σ, and 2.0 σ, respectively. Red spheres: water molecules around the inhibitors.

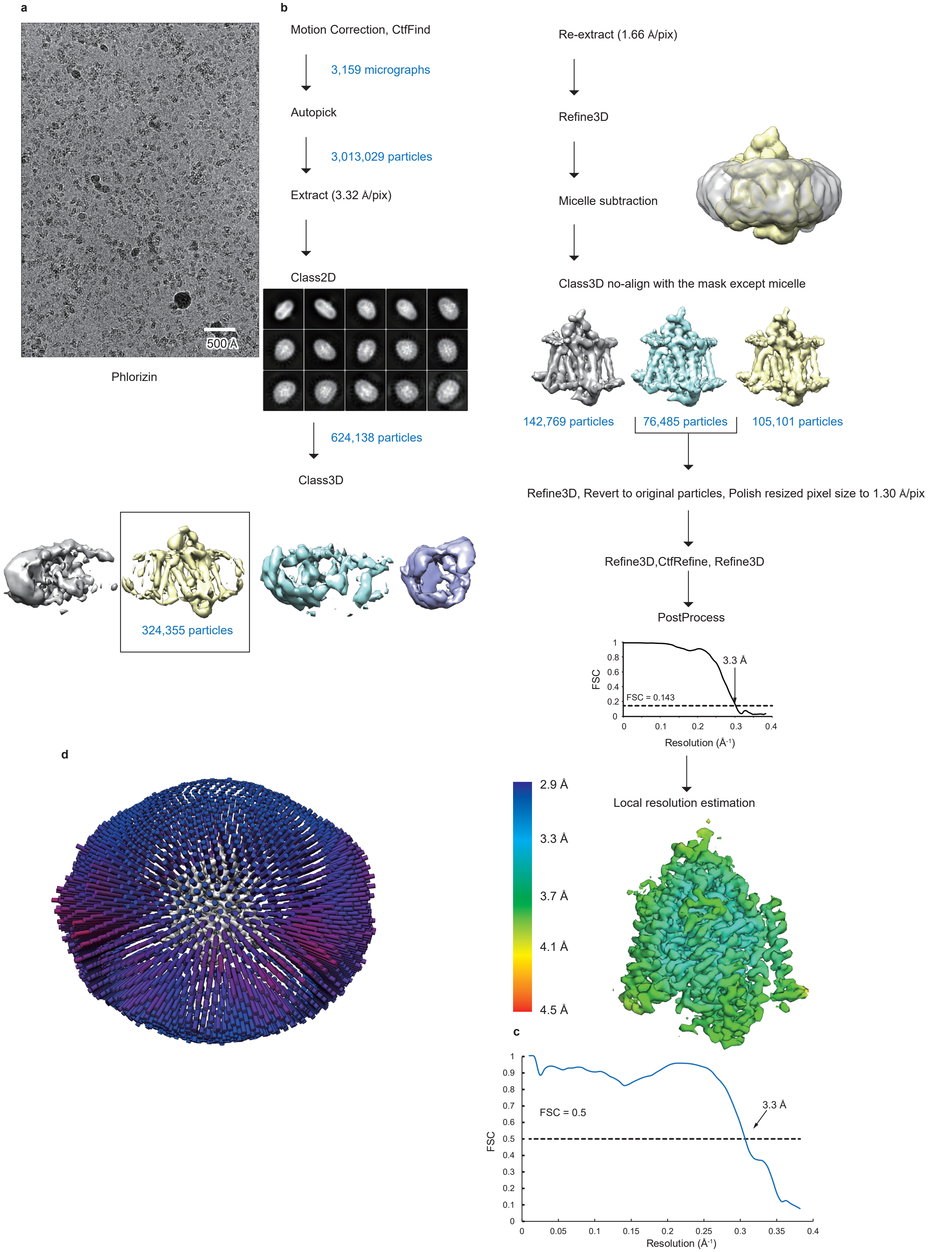


**Supplementary Figure 8. Data processing of the phlorizin-bound state.**

(a) Representative cryo-EM image of the hSGLT2–MAP17 complex in the presence of phlorizin. (b) Data processing workflow of single-particle image-processing and local-resolution analysis. Particles were separated into three groups by non-aligned 3D classification, with the mask covering the proteins (without micelles) shown in yellow. (c) Cross-validation FSC curves for map-to-model fitting. (d) Angular distributions of the final reconstruction.

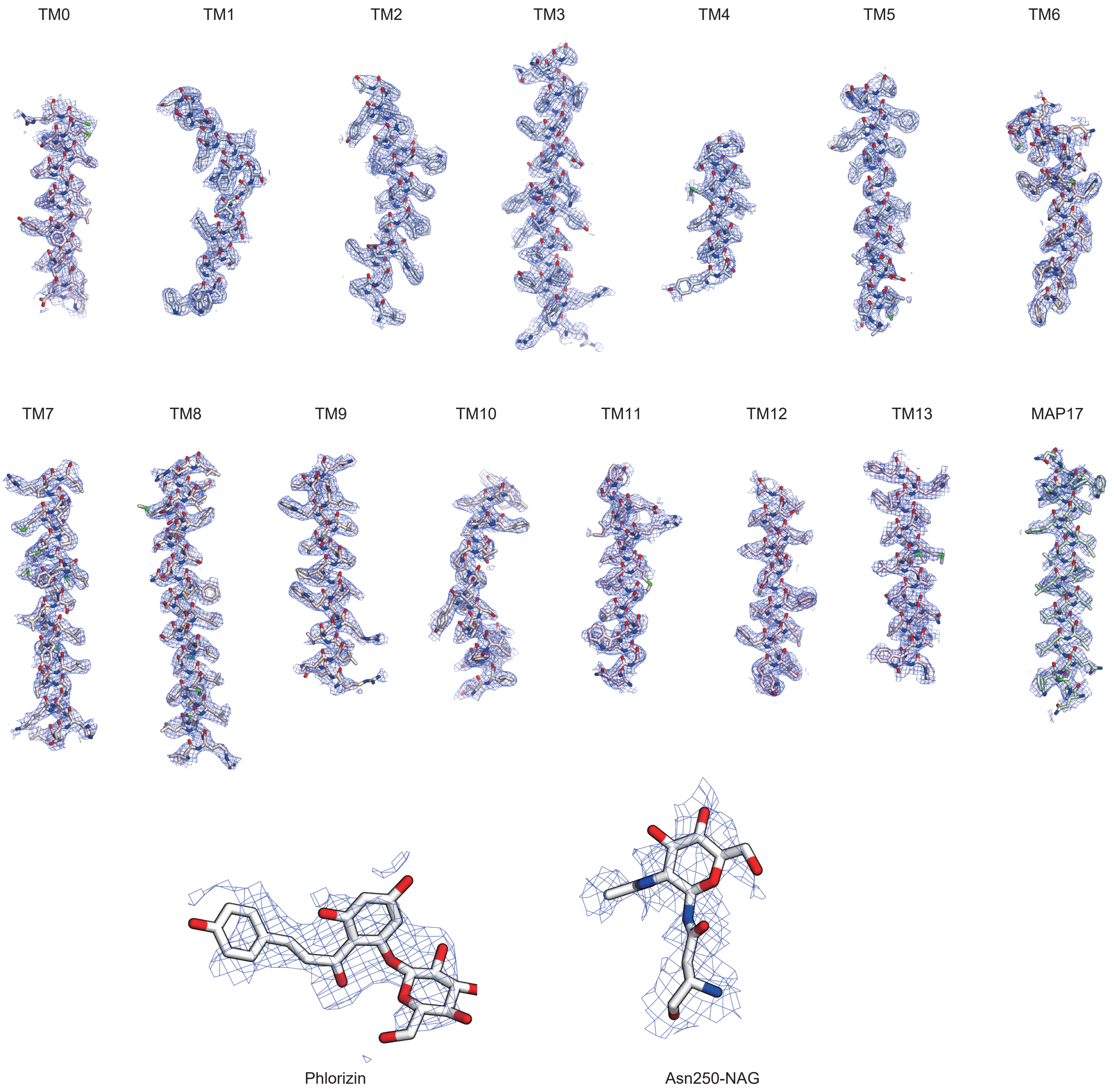


**Supplementary Figure 9. Inward-open model of hSGLT2–MAP17 in the density maps.**

The cryo-EM density and atomic models of each segment of the inward-open model of hSGLT2–MAP17, phlorizin, and glycosylation sites, contoured to 2.9 σ, 2.7 σ, and 2.7 σ, respectively.

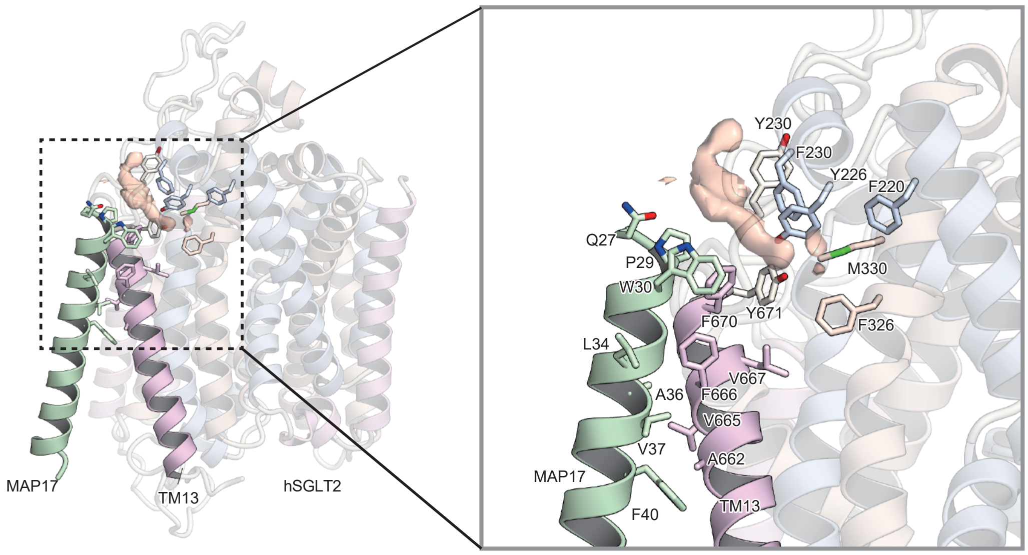


**Supplementary Figure 10. MAP17 and SGLT2 interaction site.**

The density of the lipid molecule (orange) is observed between MAP17 and SGLT2.

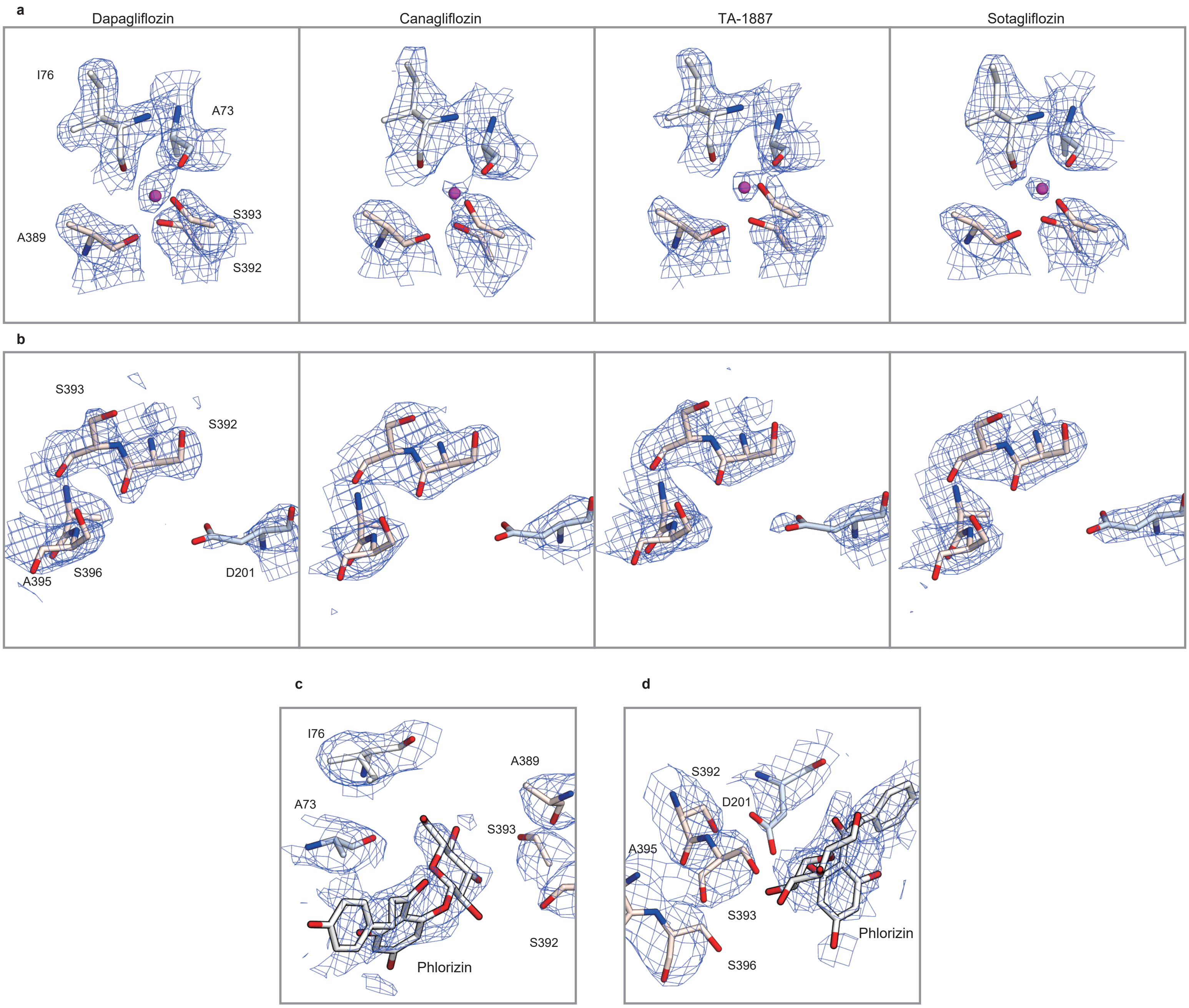


**Supplementary Figure 11. Sodium ion-binding sites of SGLT2**

(a) The sodium ion-binding Na2 sites of dapagliflozin, canagliflozin, TA-1887, and sotagliflozin are shown. (b) Sites in SGLT2 corresponding to Na3 sites where the other sodium ion binds in SGLT1. No electron density corresponding to a sodium ion can be observed. (c, d) The sodium ion-binding Na2 sites, and the sites corresponding to Na3 sites of the phlorizin-bound inward-open conformation. The phlorizin binding site is near the Na2 and Na3 sites.

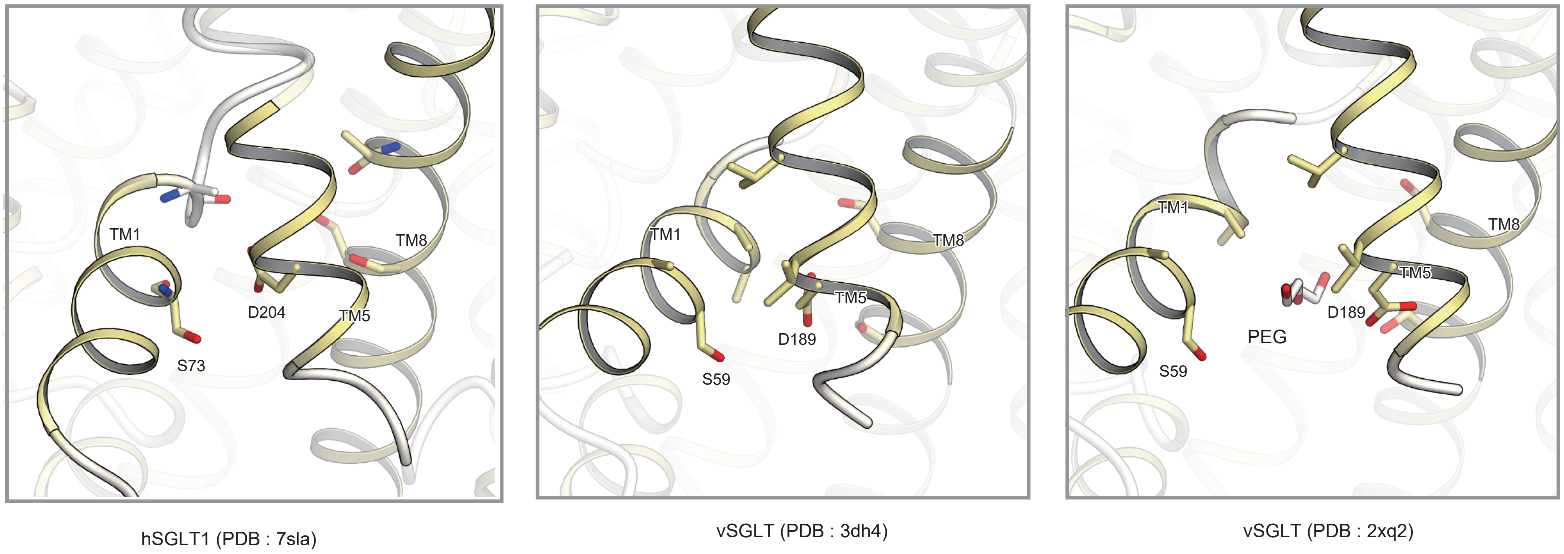


**Supplementary Figure 12. Comparison of the inward conformation of other SGLT structures.**
Structural comparison of sites corresponding to the intracellular phlorizin-binding sites of SGLT2. From left to right, inward-occluded conformation of hSGLT1, inward-occluded conformation of vSGLT, and inward-open conformation of vSGLT. The structures are viewed from the membrane side.

**Supplementary Data Table 1.**

Data collection, processing, model refinement, and validation. Clash scores, rotamer outliers, and Ramachandran plots were calculated using the Servalcat pipeline.

| **Data collection and processing** | | | | | |
|---|---|---|---|---|---|
| State | Outward | Outward | Outward | Outward | Inward |
| EMDB-ID | EMD-34673 | EMD-34705 | EMD-34610 | EMD-34737 | EMD-34823 |
| PDB ID | 8HDH | 8HEZ | 8HB0 | 8HG7 | 8HIN |
| Inhibitor | Canagliflozin | Dapagliflozin | TA-1887 | Sotagliflozin | Phlorizin |
| Microscope | Titan Krios G4 | Titan Krios G3i | | | |
| Detector | Gatan K3 Camera with Quantum LS energy filter | | | | |
| Magnification | 215,000 | 105,000 | | | |
| Voltage (kV) | 300 | | | | |
| Electron exposure ($e^{-}/Å^2$) | 64 | | | | |
| Defocus range (μm) | -0.6 to -1.6 | -0.8 to -1.6 | | | |
| Pixel size (Å/px) | 0.4 | 0.83 | | | |
| Symmetry imposed | C1 | | | | |
| Number of movies | 19,943 | 4,841 | 4,383 | 5,499 | 3,159 |
| Initial particle images | 2,364,108 | 3,692,950 | 3,395,470 | 5,242,427 | 3,013,029 |
| Final particle images | 65,919 | 197,695 | 103,853 | 72,773 | 76,485 |
| Map resolution (Å) | 3.1 | 2.8 | 2.9 | 3.1 | 3.3 |
| FSC threshold | 0.143 | 0.143 | 0.143 | 0.143 | 0.143 |
| Map sharpening B factor ($Å^2$) | -107.9 | -95.4 | -75.8 | -96.4 | -144.5 |
| **Model building and refinement** | | | | | |
| Model composition | | | | | |
| Protein atoms | 4,728 | 4,692 | 4728 | 4,763 | 4,743 |
| Metals | 1 | 1 | 1 | 1 | 0 |
| Other atoms | 49 | 46 | 49 | 44 | 45 |
| R.M.S. deviations from ideal | | | | | |
| Bond lengths (Å) | 0.011 | 0.012 | 0.012 | 0.015 | 0.018 |
| Bond angles (°) | 1.876 | 1.859 | 1.943 | 2.010 | 2.368 |
| **Validation** | | | | | |
| Clashscore | 5.62 | 6.29 | 6.66 | 6.72 | 10.28 |
| Rotamer outliers (%) | 1.20 | 2.22 | 1.20 | 2.39 | 4.58 |
| Ramachandran plot | | | | | |
| Favored (%) | 95.9 | 97.4 | 95.7 | 96.1 | 92.9 |
| Allowed (%) | 4.1 | 2.6 | 4.3 | 3.9 | 7.1 |
| Outlier (%) | 0.0 | 0.0 | 0.0 | 0.0 | 0.0 |

**Supplementary Data Table 2.**

Kinetic parameters of canagliflozin binding in the membrane fraction of hSGLT2-expressing and MAP17-expressing cells, in the presence or absence of $Na^+$.

| | Na(+) | Na(-) |
|---|---|---|
| $B_{max}$ | 99.9 ± 8.5 | 69.3 ± 6.2 |
| $K_d$ | 94.0 ± 21.7 | 109 ± 25.4 |

Data were represented in mean ± SEM (n = 3, technical replicates)

**Supplementary Data Table 3.**

Kinetic parameters of phlorizin and phloretin binding in the membrane fraction of wild-type cells or cells expressing mutated hSGLT2 and MAP17.

| Phlorizin | | | | |
|---|---|---|---|---|
| Construct | $K_{d1}$ (nM) | $B_{max1}$ (pmol/mg-protein) | $K_{d2}$ (nM) | $B_{max2}$ (pmol/mg-protein) |
| WT | 28.1 ± 11.3 | 0.356 ± 0.069 | 12400 ± 6900 | 16.8 ± 7.4 |
| S74A | 113 ± 16 | 1.16 ± 0.07 | ND | ND |
| D201 | 72.2 ± 10.0 | 0.550 ± 0.028 | ND | ND |
| F98A | ND | ND | ND | ND |
| F453A | ND | ND | ND | ND |

| Phloretin | | |
|---|---|---|
| Construct | $K_d$ (nM) | $B_{max}$ (pmol/mg-protein) |
| WT | 12000 ± 1900 | 51.7 ± 3.46 |
| S74A | ND | ND |
| D201 | ND | ND |

Data were represented in mean ± SEM (n = 3, technical replicates)
ND; not determined

**Supplementary Data Table 4.**

IC50 values of phlorizin in α-MG uptake by wild-type and mutant hSGLT2.

| Construct | IC50 (nM) |
|---|---|
| WT | 57.0 ± 8.49 |
| S74A | 145 ± 27 |
| F98A | 1640 ± 1720 |
| F453A | 468 ± 200 |

Data were represented in mean ± SEM (n = 4, biological replicates)
